# SNTA1 Gene Rescues Ion Channel Function in Cardiomyocytes Derived from Induced Pluripotent Stem Cells Reprogrammed from Muscular Dystrophy Patients with Arrhythmias

**DOI:** 10.1101/2022.01.25.477696

**Authors:** Eric N Jimenez-Vazquez, Michael Arad, Álvaro Macías, Maria Linarejos Vera-Pedrosa, Francisco M. Cruz-Uréndez, Ashley J Cuttitta, André Monteiro Da Rocha, Todd J Herron, Daniela Ponce-Balbuena, Guadalupe Guerrero-Serna, Ofer Binah, Daniel E Michele, José Jalife

**Author notes:** Correspondence to José Jalife, MD, PhD: Centro Nacional de Investigaciones Cardiovasculares (CNIC) Carlos III, Melchor Fernández Almagro 3, 28029, Madrid, SPAIN.

## Abstract

Patients with cardiomyopathy of Duchenne Muscular Dystrophy (DMD) are at risk of developing life-threatening arrhythmias, but the mechanisms are unknown. We aimed to determine the role of cardiac ion channels controlling cardiac excitability in the mechanisms of arrhythmias in DMD patients. To test whether cardiac dystrophin mutations lead to defective Na_V_1.5–Kir2.1 channelosomes and arrhythmias, we generated iPSC-CMs from two hemizygous DMD males, a heterozygous female, and two unrelated controls. Two Patients had abnormal ECGs with frequent runs of ventricular tachycardia. iPSC-CMs from all DMD patients showed abnormal action potential profiles, slowed conduction velocities, and reduced sodium (*I*_Na_) and inward rectifier potassium (*I*_K1_) currents. Membrane Na_V_1.5 and Kir2.1 protein levels were reduced in hemizygous DMD iPSC-CMs but not in heterozygous iPSC-CMs. Remarkably, transfecting just one component of the dystrophin protein complex (α1-syntrophin) in hemizygous iPSC-CMs restored channelosome function, *I*_Na_ and *I*_K1_ densities and action potential profile. We provide the first demonstration that iPSC-CMs reprogrammed from skin fibroblasts of DMD patients with cardiomyopathy have a dysfunction of the Na_V_1.5-Kir2.1 channelosome, with consequent reduction of cardiac excitability and conduction. Altogether, iPSC-CMs from patients with DMD cardiomyopathy have a Na_V_1.5-Kir2.1 channelosome dysfunction, which can be rescued by the scaffolding protein α1-syntrophin to restore excitability.

## 1. Introduction

Null mutations in the Dp427 isoform of the dystrophin gene result Duchenne Muscular Dystrophy (DMD).^1^ This inheritable X-linked disease affects primarily adolescent males causing progressive skeletal muscle deterioration, with negative effects in the central nervous system.^2^ Muscular dystrophies are also characterized by cardiac muscle involvement,^3^ which usually starts with an abnormal ECG.^4^ Eventually, most patients with DMD will develop cardiomyopathy by 20 years of age.^5^ Many will be at a high risk for arrhythmia and sudden cardiac death (SCD), which contributes considerably to the morbidity and mortality of the disease.^6^ However, diagnosis and prevention of arrhythmia is challenging in DMD patients.^7^

The mechanisms responsible for arrhythmias and SCD in patients with DMD cardiomyopathy are poorly understood. The dystrophin associated protein complex (DAPC) is involved in mechanoprotection of the plasma membrane.^8^ The DAPC acts also as a putative cellular signaling complex that forms a scaffold for numerous signaling and membrane ion channel proteins.^9–11^ Absence of dystrophin in DMD has the potential to alter trafficking, localization and function of DAPC associated proteins in skeletal and cardiac muscle.^12^ For example, the expression and function of ion channels are defective in ventricular cardiomyocytes of the *mdx* mouse model.^10, 13–16^ Absence of dystrophin in young *mdx* mice affects the function of Na_V_1.5, leading to cardiac conduction defects.^10^ Inward rectifier potassium current *I*_K1_ is reduced in the *mdx* mouse^14^ but the consequences of the disruption have not been identified.

Results from our laboratory and others strongly suggest that Na_V_1.5 and Kir2.1 control cardiac excitability by mutually modulating each otheŕs surface expression.^11, 16–20^ At the lateral membrane, Na_V_1.5 and Kir2.1 channels form macromolecular complexes (“channelosomes”)^21^ that include α1-syntrophin, which is a part of the DAPC.^10^ Thus, we hypothesize that dystrophin gene mutations that truncate the Dp427 dystrophin isoform, disrupt Na_V_1.5 - α1-syntrophin - Kir2.1 interactions, altering the function of the most important ion channels controlling cardiac excitability and conduction velocity, which would place the DMD patient at risk of arrhythmogenesis and SCD.

Here we have used matured ventricular-like iPSC-CMs derived from two genetically distinct hemizygous DMD males, a heterozygous DMD female and two unrelated healthy subjects (controls) to investigate the mechanisms underlying the arrhythmias associated with loss-of-function dystrophin mutations. We demonstrate that iPSC-CMs from patients with DMD cardiomyopathy have a dysfunction of the Na_V_1.5-Kir2.1 channelosome, which can be rescued by transfection with *SNTA1*, the gene coding the DAPC-related scaffolding protein α1-syntrophin.

## 2. Methods

**(See Supplemental Methods for details)**

### 2.1 Ethics statement

We obtained skin biopsies from 2 hemizygous DMD patients, 1 heterozygous female, and 2 healthy subjects after written informed consent in accordance with the Helsinki Committee for Experiments on Human Subjects at Sheba Medical Center, Ramat Gan, Israel (Approval number: 7603-09-SMC), and with IRB HUM00030934 approved by the University of Michigan Human IRB Committee. The use of iPS cells and iPSC-CMs was approved by the Human Pluripotent Stem Cell Research Oversight (HPSCRO, #1062) Committee of the University of Michigan and the Spanish National Center for Cardiovascular Research (CNIC) Ethics Committee and the Regional Government of Madrid. Data will be available upon rationale request.

### 2.2 Generation of iPSCs

Cell lines were generated using Sendai virus CytoTune-iPS 2.0 Sendai reprogramming kit (Thermo Fisher) for transfection of Yamanaka’s factors, as described.^22, 23^

### 2.3 Patient-specific iPSC-CMs monolayers (adapted from Herron et al. 2016).^24^

We obtained highly purified iPSC-CMs after directed cardiac differentiation. After 30 days in culture, cardiomyocytes were purified, dissociated and plated on Matrigel-coated polydimethylsiloxane (PDMS) membranes at a density of ∼200K cells per monolayer. Cells were maintained for 7 days before re-plating onto Matrigel-coated micropatterned PDMS for patch-clamp and immunostaining experiments. At least 3 separate cardiomyocyte differentiations were used for all the experiments.

### 2.4 Micropatterning on PDMS (adapted from ref^25^)

Stamps were sonicated and then incubated with Matrigel diluted in water (Corning, 100 μg/mL) for 1 h. Then, 18 mm PDMS circles were UVO treated before micropatterning. An hour later, the Matrigel solution from the PDMS stamps was aspirated and each stamp was inverted onto each PDMS circle and removed one by one. The micropatterned PDMS was incubated overnight with pluronic-F127 at room temperature. Then, it was cleaned with antibiotic-antimycotic solution and exposed to UV light before re-plating cells. About 30,000 human iPSC-CMs were placed in the center of the micropatterned area. Cells were cultured on micropatterns at least 4 days prior to experiments.

### 2.5 Electrophysiology

We used standard patch-clamp recording techniques to measure the action potentials (APs), as well as sodium current (*I*_Na_), L-type calcium current (*I*_CaL_), and inward rectifier potassium current (*I*_K1_) in the whole-cell configuration. All experiments were conducted at room temperature, except for the AP recordings, which were obtained at 37°C and paced at 1 and 2 Hz.

### 2.6 RT-PCR

For quantitative evaluation of mRNA expression in each experimental group, total RNA was prepared using the RNeasy Mini Kit (Qiagen), including DNAse treatment. cDNA was synthetized using SuperScript III First-Strand Synthesis System (Invitrogen). Quantitative PCR was performed using TaqMan Universal PCR Master Mix (Applied Biosystems) in the presence of primers for *SCN5A*, *CACNA1C* and *KCNJ2*. We calculated mRNA fold expression by the ΔΔCT method using the 18S rRNA as the housekeeping gene. Every qPCR reaction was performed in triplicate and repeated using cDNA from at least 3 separate cardiomyocyte differentiation cultures.

### 2.7 Western Blotting

Standard Western blotting was applied and Image Lab software (Bio-Rad) was used for analysis. Total and biotinylated protein was obtained from iPSC-CM monolayers and resolved on SDS-PAGE gels. Membranes were probed with anti-human Dystrophin, Na_V_1.5 and Kir2.1 antibodies, using Actinin as the loading control for total protein analysis, Na/K-ATPase for biotinylation experiments, and cTnT as the marker for cardiomyocytes.

### 2.8 Immunofluorescence

iPSC-CMs were plated on micropatterned PDMS, fixed, treated and analyzed as described in detail in *Supplemental Methods* (see also ref^24^) Images were recorded with a Nikon A1R confocal microscope (Nikon Instruments Inc.) and Leica SP8 confocal microscope (Leica Microsystems).

### 2.9 Optical Mapping

Optical action potentials were recorded using the voltage-sensitive fluorescent dye FluoVolt (F10488; Thermo Scientific). Activation patterns were determined, and conduction velocity was measured as described previously.^24, 26^

### 2.10 Generation and Stable Transfection of *SNTA1-IRES-GFP*

Non-viral piggy-bac vector encoding SNTA1-IRES-GFP were co-transfected with mouse transposase-expression vector into iPSCs cells. After 3–5 days GFP positive cells were selected by FACS sorter and grow-up. Every week, fluorescence was confirmed, and cells sorted to confirm cDNA stable integration into the cells. After that, iPSC-CMs differentiation protocol was applied as stated above.

### 3.0 Statistics

All data are expressed as mean ± SEM. In each data set a Grubbs’ test was performed after data collection to determine whether a value should be considered as a significant outlier from the rest. Nonparametric Mann-Whitney test was used. Multiple comparisons were tested using two-way analysis of variance (ANOVA) followed by *Sidak’s* or Dunnett’s test using Prism 8. *P* < 0.05 was considered significant. All experiments were performed as a single-blind study to avoid sources of bias.

## 3. Results

### 3.1 Clinical characteristics

We generated iPSC-CM lines from reprogrammed skin fibroblasts that were collected from 3 patients suffering from DMD cardiomyopathy. Two male hemizygous had a clinical and genetic diagnosis for DMD; the third patient was a DMD heterozygous female (*Figure 1*). iPSC-CMs from a healthy subject unrelated to the patients (Control 1) and a line of BJ iPSC-CMs (Control 2) acted as negative controls. Complete clinical data were accessible for one DMD male and the heterozygous female. The hemizygous male (Male 1) harboring a nonsense point mutation in the dystrophin gene (exon 41) experienced DMD from early childhood, being diagnosed with dilated cardiomyopathy at age 17. Eight years later he was hospitalized in respiratory and heart failure (LVEF = 15%), requiring tracheostomy and prolonged ventilation. An ECG exhibited sinus rhythm with a narrow QRS and QR pattern in L1, AVL and QS leads V2*–*3 (*Figure 1a*). At age 30, the patient became respirator-dependent with a reasonably controlled heart failure. A routine Holter-ECG obtained three years later showed frequent premature ventricular complexes and episodes of non-sustained ventricular tachycardia at rates of up to 200/min. An ICD was implanted, which discharged appropriately 2 years later for repeated episodes of ventricular flutter deteriorating into ventricular fibrillation (*Figure 1b*). Three years later, the patient expired of heart failure at age 38.

**Figure 1.**
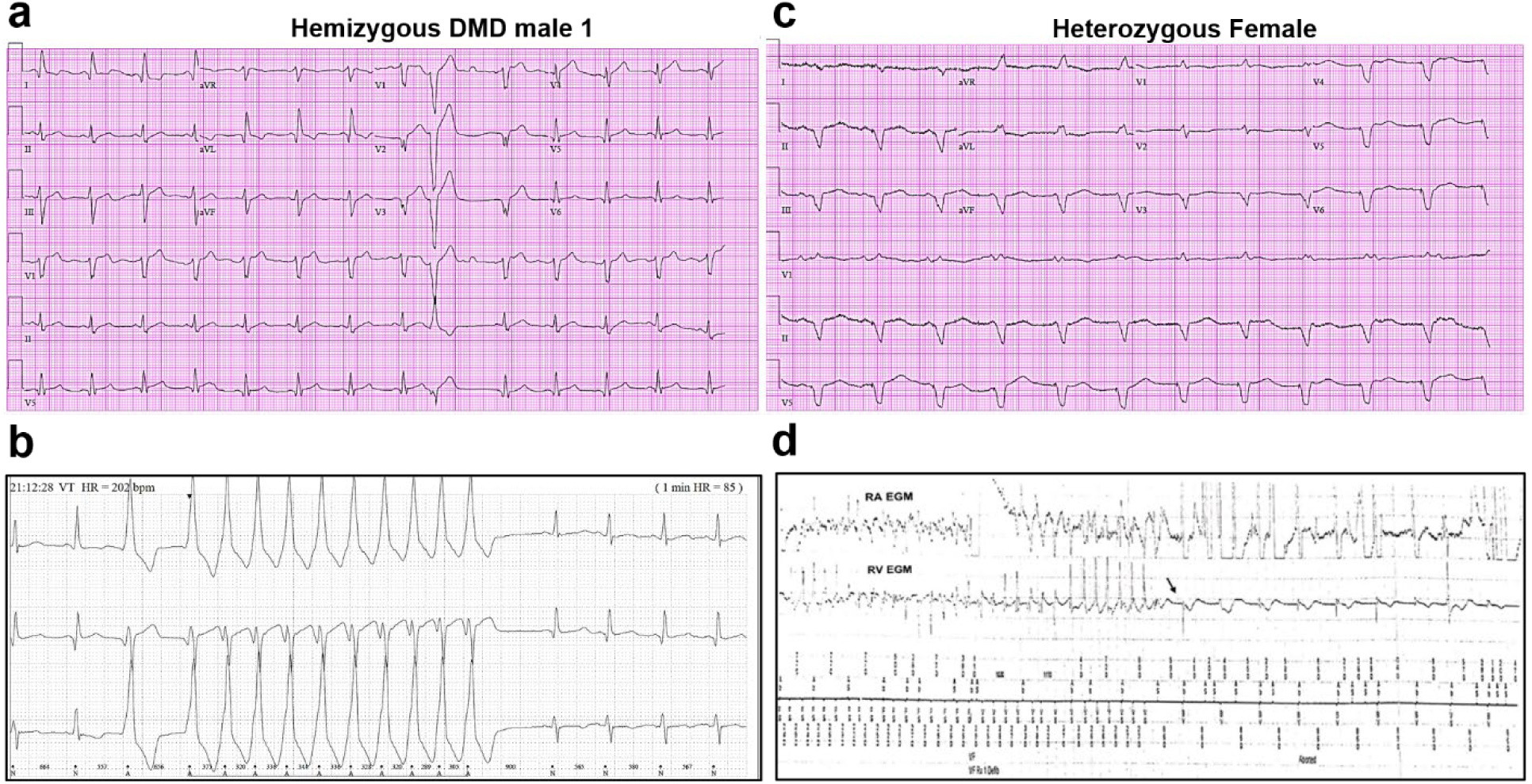
Altered ECG and arrhythmias in DMD patients with cardiomyopathy. **(a)** Abnormal ECG in a 34-year-old DMD male: PR interval, 116 ms; QRS, 120 ms; QT/QTc, 404/472 ms; and PRT axes, 18-16-90. **(b)** Holter recording from the same patient shows non-sustained monomorphic ventricular tachycardia. **(c)** Abnormal ECG from the heterozygous female at 50-years of age: left axis deviation; QRS, 178 ms; QT/QTc, 564/612 ms; and PRT axes, 55-263-85. **(d)** Holter atrial electrograms of the heterozygous female shows atrial fibrillation with complete AV block after AV nodal ablation. Ventricular electrogram shows polymorphic ventricular tachycardia with spontaneous termination (arrow) and resumption of ventricular pacing.

The female patient, heterozygous for a deletion of 5 exons (Δ8–12) in the dystrophin gene, presented proximal muscle weakness with creatine kinase elevation at age 42. She had a son with DMD who died at 16. At presentation, she exhibited biventricular dysfunction with left ventricular dimension of 65 mm, LVEF of 30% and moderate to severe mitral insufficiency. At age 49, she developed severe biventricular dysfunction with LVEF=20% and severe tricuspid regurgitation. She was in NYHA IV, and the cardio-respiratory exercise test showed a VO_2_ max of 6 mL/kg/min, indicating a severely reduced aerobic capacity. An ECG obtained at age 50 revealed severe QRS widening and QT prolongation (*Figure 1c*). At that time, she had LVEF 30–35% and her heart failure was relatively well controlled. A year later, she developed paroxysmal atrial fibrillation with rapid ventricular response and recurrent episodes of non-sustained ventricular tachycardia (*Figure 1d*). AV nodal ablation and CRTD pacemaker-defibrillator implantation was required. The patient died at 51 in end-stage heart failure associated with renal insufficiency.

The additional DMD patient (Male 2) was a 13-year-old male carrying a 6-exon dystrophin deletion (Δ45–50). The patient was non-ambulatory (used a motorized wheelchair) but respirator free at the time of the skin biopsy. He did not have significant cardiomyopathy at the time of collection, which was not surprising given his young age and the typical presentation of DMD cardiomyopathy as later onset.^27, 28^ No follow-up information is available for this patient. The unrelated healthy individuals (Controls 1 and 2) have no personal or family history of DMD or any related disease.

### 3.2 Dystrophin is absent in iPSC-CMs derived from hemizygous DMD patients

Compared to Control-1 iPSC-CMs and to left ventricle samples from a patient with Becker dystrophy, iPSC-CMs from hemizygous males were deficient in the full-length adult DP427 dystrophin isoform (*Figure 2a–b*). The iPSC line named Male 2 shows a deletion of exons 45–50, while the other dystrophic cell line (Male 1) presents a nonsense point mutation (R1967X) in exon 41 of the dystrophin gene constituting a premature stop codon. The cell line generated from the 50-year-old DMD heterozygous female carried a deletion of exons 8–12. Notably, her iPSC-CMs showed expression of dystrophin protein like the control (*Figure 2a–b*).

**Figure 2.**
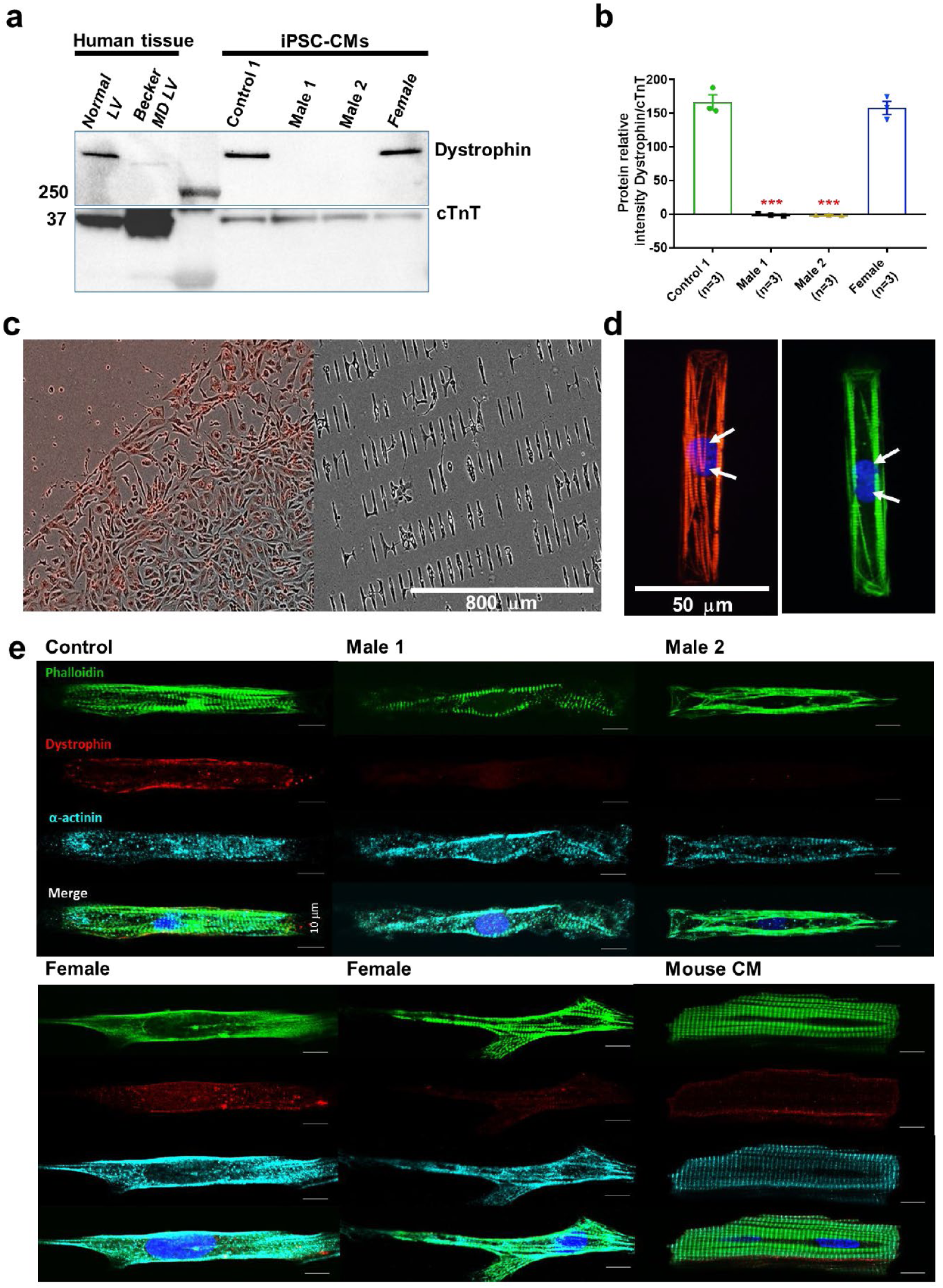
DMD patient-specific iPSC-CMs do not express dystrophin. **(a)** Top right, control and heterozygous female iPSC-CMs express dystrophin. iPSC-CMs from hemizygous dystrophic cell lines (Male 1 and Male 2) did not express the large dystrophin isoform. Top left, control tissue lysates from a normal individual and a patient with Becker MD. Dystrophic left ventricular tissue did express dystrophin, but to a lesser extent than normal left ventricle tissue. These data were generously provided by the Hypertrophic Cardiomyopathy Clinic, University of Michigan. **(b)** Quantitation of dystrophin in control and heterozygous female iPSC-CMs. Dystrophin was absent in DMD iPSC-CMs (\*\*\**P* = 0.0001) compared to control iPSC-CMs. Heterozygous female cells exhibited nearly normal dystrophin expression (*P* = 0.5864). Protein concentration confirmed by Western blot against troponin T. Two-tailed Mann-Whitney test. Errors bars, SEM. The *n*-values are in parentheses. **(c–e)** iPSC-CMs plated onto Matrigel-coated micropatterned PDMS. **(c)** Male 1 iPSC-CMs plated as a monolayer on a Matrigel-coated PDMS (left) for 1 week, and then dissociated for re-plating onto micropatterned PDMS (right). **(d)** Control iPSC-CMs fixed and stained on micropatterns. Immunostaining for cardiac troponin I (red) and F-actin (green). Nuclei were stained with DAPI (white arrows). Scale bar, 50 µm**. (e)** Immunostaining for dystrophin in iPSC-CMs from control, dystrophic Male 1 and Male 2, female, and mature mouse cardiomyocytes. DMD cells do not express dystrophin compared to control. Heterozygous female iPSC-CMs showed variable expression of dystrophin. Scale bar, 10 µm.

### 3.3 Micropatterning controls cell shape and facilitates electrophysiological recordings

Cell shape is critical for cardiomyocyte electrical, mechanical and contractile function.^25^ Adopting the typical cylindrical morphology helps improve contractility, which promotes electrophysiological phenotype maturation.^29^ When cultured on a non-micropatterned smooth surface, DMD iPSC-CMs are flat-shaped and have a frail membrane making them a challenge for patch-clamp experiments (*Figure 2c, left*). Therefore, we plated and fixed our iPSC-CMs on Matrigel-coated micropatterned PDMS (*Figure 2c, right*). The approach produces large numbers of thick cylindrical-shaped, binucleated cardiomyocytes with well-organized sarcomeres (*Figure 2d*), which are two important signs of maturation. Micropatterned iPSC-CMs are easier to patch. They are electrically excitable and their electrical phenotype approaches the adult human cardiomyocyte, with maximum diastolic potentials (MDP) of -70 to -80 mV, and action potential durations (APDs) of 200–300 ms (see below).^30, 31^ On the other hand, as shown in *Figure 2e*, unlike control cells, immunostained DMD cells do not express dystrophin, whereas iPSC-CMs from the female patient show variable expression of dystrophin.

### 3.4 Action potentials in dystrophic iPSC-CMs have a reduced maximum upstroke velocity

Clinically, DMD patients may experience cardiac complications and often exhibit electrical conduction abnormalities and life-threatening arrhythmias (see *Figure 1* above).^32, 33^ At the cellular level, such alterations are often the result of reduced excitability. We therefore conducted patch-clamp recordings in micropatterned iPSC-CMs in the current-clamp configuration. In *Suppl. Tables 1–3* we present comparisons at two different frequencies for DMD versus Control 1 (*Suppl. Table 1*), DMD versus Control 2 (*Suppl. Table 2*) and Control 1 vs Control 2 (*Suppl. Table 3*). We quantitated AP parameters such as maximal upstroke velocity (dV/dt_max_), overshoot, AP amplitude, MDP and AP duration. Statistical analysis demonstrated that Control 1 and Control 2 were very similar to each other, both exhibiting well-polarized MDPs, dV/dt_max_ larger than 40 V/sec and amplitudes larger than 100 mV. However, they both differed significantly from all three DMD groups (*see Figure 3 and Suppl. Figure 1a–f*), particularly in terms of dV/dt_max_. iPSC-CMs from both DMD male and female patients revealed abnormal AP profiles compared to both controls. For example, Overshoot and amplitude were lower in the Male 2 cells compared to the controls. In addition, female DMD cells showed a more depolarized MDP than control iPSC-CMs (*Figure 3e*). Finally, no significant differences existed in APD_90_ values and similar action potential parameter changes were obtained at 2 Hz (*Suppl. Table 1, Suppl. Figures 1a–f* and *2*).

**Figure 3.**
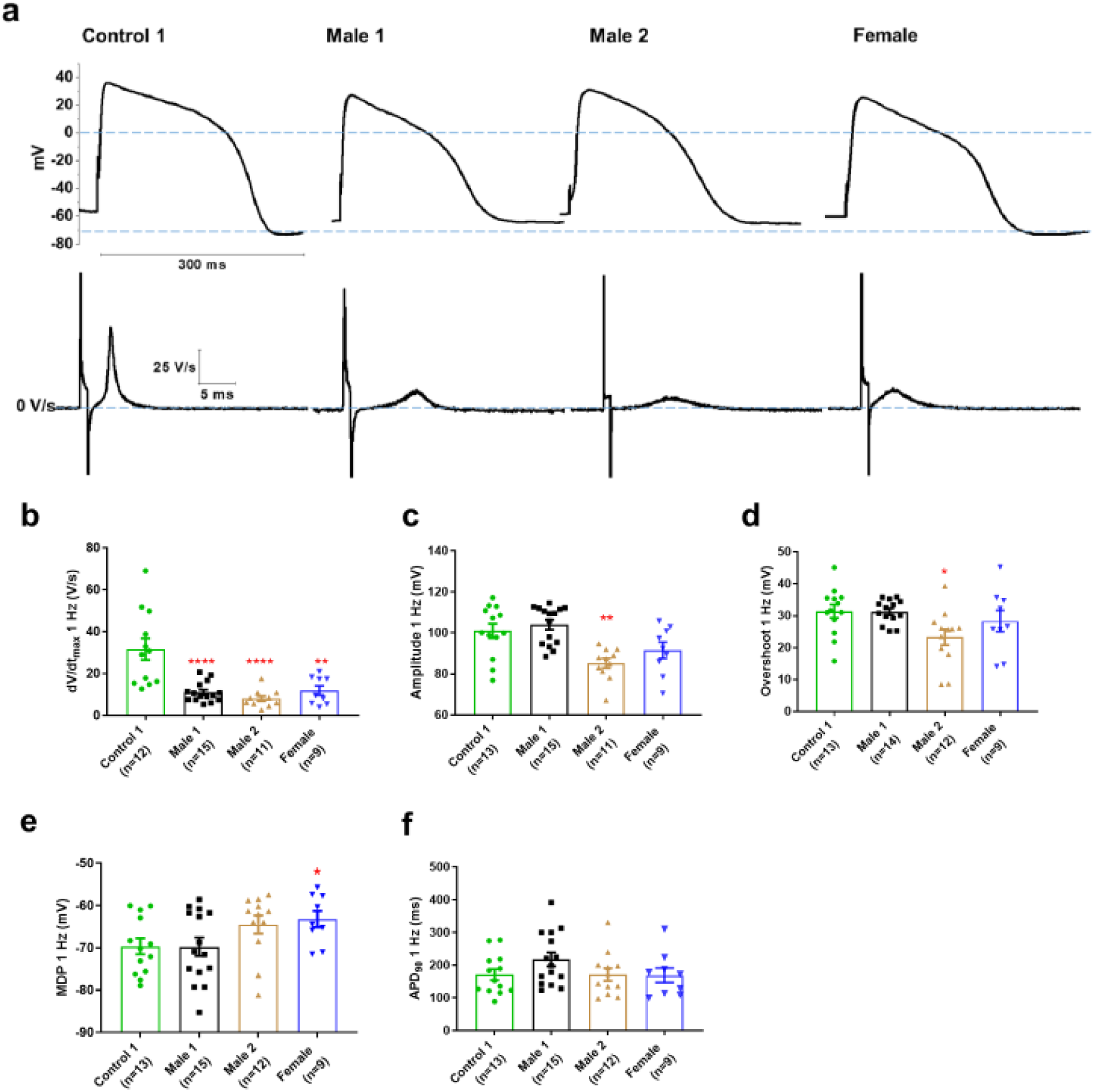
Action potential properties in control, DMD, and female iPSC-CMs. **(a)** Representative action potentials of ventricular-like iPSC-cardiomyocytes from Control 1, heterozygous female and DMD individuals. The respective dV/dt trace is shown below each action potential. **(b)** Mann-Whitney test revealed that dV/dt_max_ was reduced in both DMD compared to Control 1. dV/dt_max_ was also significantly reduced in the female cells. **(c–e)** Overshoot and Amplitude were only affected in the Male 2 iPSC-CMs, while heterozygous female cells were significantly more depolarized compared to control. **(f)** APD_90_ was similar in all iPSC-CMs tested. Cells plated on micropatterns were paced at 1 Hz. Errors bars, SEM. The *n*-values are in parentheses. Two-tailed Mann-Whitney test. \*\*\*\**P* = 0.0001 and \**P* < 0.05.

### 3.5 Conduction velocity is impaired in DMD iPSC-CM monolayers

The reduced dV/dt_max_ at the single cell level suggested that conduction velocity (CV) may be compromised in iPSC-CMs monolayers from affected individuals. Hence, we conducted optical mapping experiments using the voltage-sensitive fluorescent dye FluoVolt^TM^ in control, DMD, and female iPSC-CM monolayers paced at various frequencies (*Figure 4a*). CV in dystrophin-deficient iPSC-CM monolayers was 50% slower than control monolayers paced at 1 Hz (27 ± 2 cm/s and 29 ± 4 cm/s in hemizygous Male 1 and Male 2 cells, respectively, versus 56 ± 3 cm/s in control cells, *Figure 4b–c*). CV of Control 2 monolayers was 42 ± 5 cm/s (*Suppl. Figure 3*). Remarkably, CV in the heterozygous female monolayers was even slower (18 ± 3 cm/s). In all three groups, the CV restitution curve displayed slightly slower velocities at higher frequencies (*Figure 4d*). Most important, in the female monolayer (*Figure 4e*), slower and more heterogeneous patterns of electrical wave propagation were accompanied by focal discharges in the form of trigeminy (*Figure 4e*, *left*), which often triggered unidirectional block and reentry (*Figure 4e, right*, and *Suppl. Video 1* and *2*). Altogether, the data presented in *Figures* 3 and 4 provide a direct mechanistic explanation for the conduction abnormalities and arrhythmias seen on the ECGs of at least two of the patients (see *Figure 1*). In all three iPSC-CMs from affected individuals, the reduced CV occurred in the absence of measurable changes in connexin43 (Cx43) protein (*Suppl. Figure 4*). We did not detect any significant differences in Cx43 expression among control, heterozygous, and hemizygous iPSC-CMs in these monolayer experiments.

**Figure 4.**
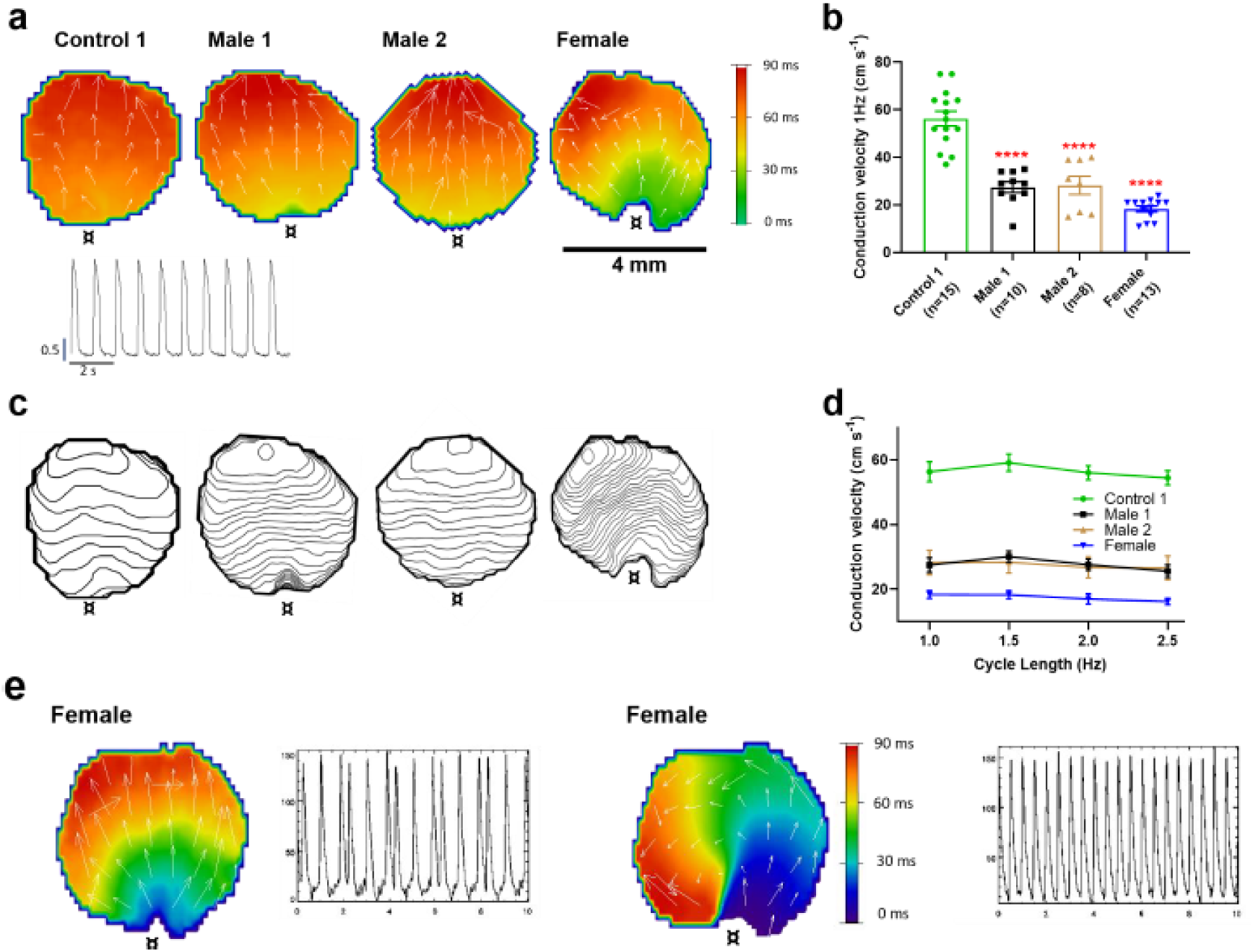
Conduction velocity is slower in iPSC-CM monolayers from DMD hemizygous male and heterozygous female than control. **(a)** Activation maps of action potential propagation at 1 Hz. Each color represents a different activation time with time zero appearing in green (**¤** indicates the location of the stimuli for each monolayer). White vectors (↑) are a measure of local velocity and direction of the wave. *Inset.* Representative optical APs at 1 Hz. **(b)** Bar graphs of CV in each monolayer group, as indicated. Numbers in parenthesis are number of monolayers per group. **(c)** Averaged 2-ms contour isochrone maps for each representative monolayer above. Tighter averaged isochrone contours in the hemizygous and heterozygous iPSC-CM monolayers indicate slowed and more heterogeneous CV compared to control. **(d)** CV restitution tended to slow in all groups as pacing frequency increased. **(e)** Arrhythmias in heterozygous female iPSC-CMs monolayers (see also *Suppl. Videos*). Left map, spontaneous pacemaker activity; *Left inset*, single pixel recording reveals premature ectopic discharges in a pattern of trigeminy; Right map, high-frequency reentrant tachycardia maintained by a self-sustaining rotor; *Right inset*, single pixel recording shows the interbeat interval (500 ms) of the reentrant tachycardia. Errors bars represent SEM. The *n*-values are in parentheses. Two-tailed Mann-Whitney test. \*\*\*\**P* < 0.0001.

### 3.6 Sodium current is down-regulated in DMD iPSC-CMs

Sodium channels determine the upstroke velocity of the cardiac action potential and consequently play a key role in the conduction of the cardiac electrical impulse.^34^ Here we compared the sodium current (*I*_Na_) density in the DMD male and female iPSC-CMs versus each of the controls. In *Figure 5a* and *b*, the peak inward *I*_Na_ density in hemizygous iPSC-CMs was significantly decreased (-14 ± 1 pA/pF for Male 1 cells and -15 ± 1 pA/pF for Male 2 cells) compared to both Control 1 (-27 ± 3 pA/pF) and Control 2 iPSC-CMs (-38 ± 1 pA/pF; *Suppl. Figure 1g*, *Suppl. Table 4* and *5*). Importantly, the *I*_Na_ density in heterozygous female cells was also dramatically reduced (-11 ± 1 pA/pF). Altogether, except for peak sodium current density, statistical comparisons in terms of biophysical properties of I_Na_ (half maximal activation, slope factor, reversal potential) for DMD vs Control 1 (*Suppl. Table 4*), DMD vs Control 2 (*Suppl. Table 5*) and Control 1 vs Control 2 (*Suppl. Table 6*) showed no differences among any of the groups. Also, as shown in *Suppl. Figure 5*, cell capacitance in all the patient-specific cells was similar to control, indicating that cell size was similar in all groups.

**Figure 5.**
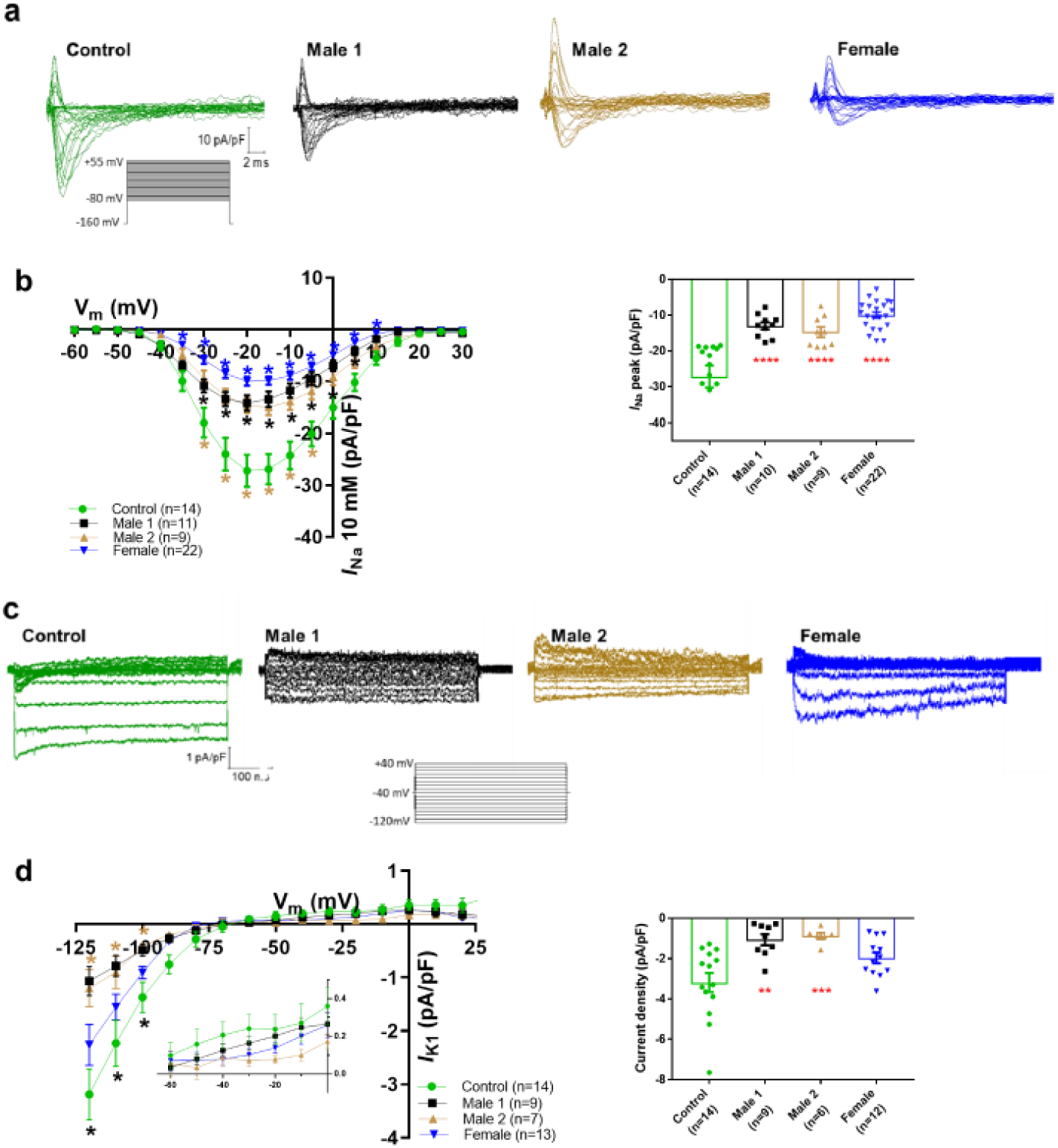
Sodium (*I*_Na_) and Inward rectifier potassium (*I*_K1_) channel properties in control, DMD, and female iPSC-cardiomyocytes. **(a)** Superimposed *I*_Na_ current traces for Control 1, hemizygous and heterozygous iPSC-CMs elicited by the pulse protocol shown by the inset. **(b) Left**, normalized current-voltage (I/V) relationships. *I*_Na_ was significantly reduced in both Male 1 and Male 2 iPSC-CMs compared with control at the specified voltages. Heterozygous female iPSC-CMs showed also a very reduced current density from 35 to 10 mV. Two-way analysis of variance (ANOVA) followed by Sidak’s multiple comparisons test. **Right,** peak *I*_Na_ density at 20 mV was reduced in all three affected groups compared to control. **(c)** Typical *I*_K1_ density traces from control and DMD cells elicited by the pulse protocol in the *inset*. **(d) Left**, I/V relationships. *I*_K1_ was significantly reduced in both Male 1 and Male 2 iPSC-CMs compared with control at the specified voltages. Two-way ANOVA followed by Sidak’s multiple comparisons. **Right,** normalized current densities at -120 mV. *I*_K1_ was decreased in Male 1 and in Male 2 cells compared to control cells. Two-tailed Mann-Whitney test. Errors bars represent SEM. The *n*-values are in parentheses. ****P < 0.0001, \*\**P* < 0.005 and \**P* < 0.05 and *P < 0.056.

The above data indicate that dystrophin deficiency reduces the *I*_Na_ density, which may be considered one of the main causes for the cardiac conduction defects reported in DMD patients.^35, 36^ The absence of dystrophin might also affect other ionic currents. For instance, the L-type calcium current (*I*_Ca,L_) is increased in cardiomyocytes from adult *mdx* mice.^13, 37^ In addition, as previously suggested, *I*Ca,L density is increased in iPSC-CMs from DMD patients.^35^ However, under our experimental conditions, *I*_Ca,L_ was unaltered in hemizygous and heterozygous DMD iPSC-CMs (*Suppl. Tables 4–6* and *Suppl. Figures 1h* and *6*). Differences in culture conditions and cell maturation (see Methods and Ref^24^) might have contributed to the different outcomes in the two studies.

**Figure 6.**
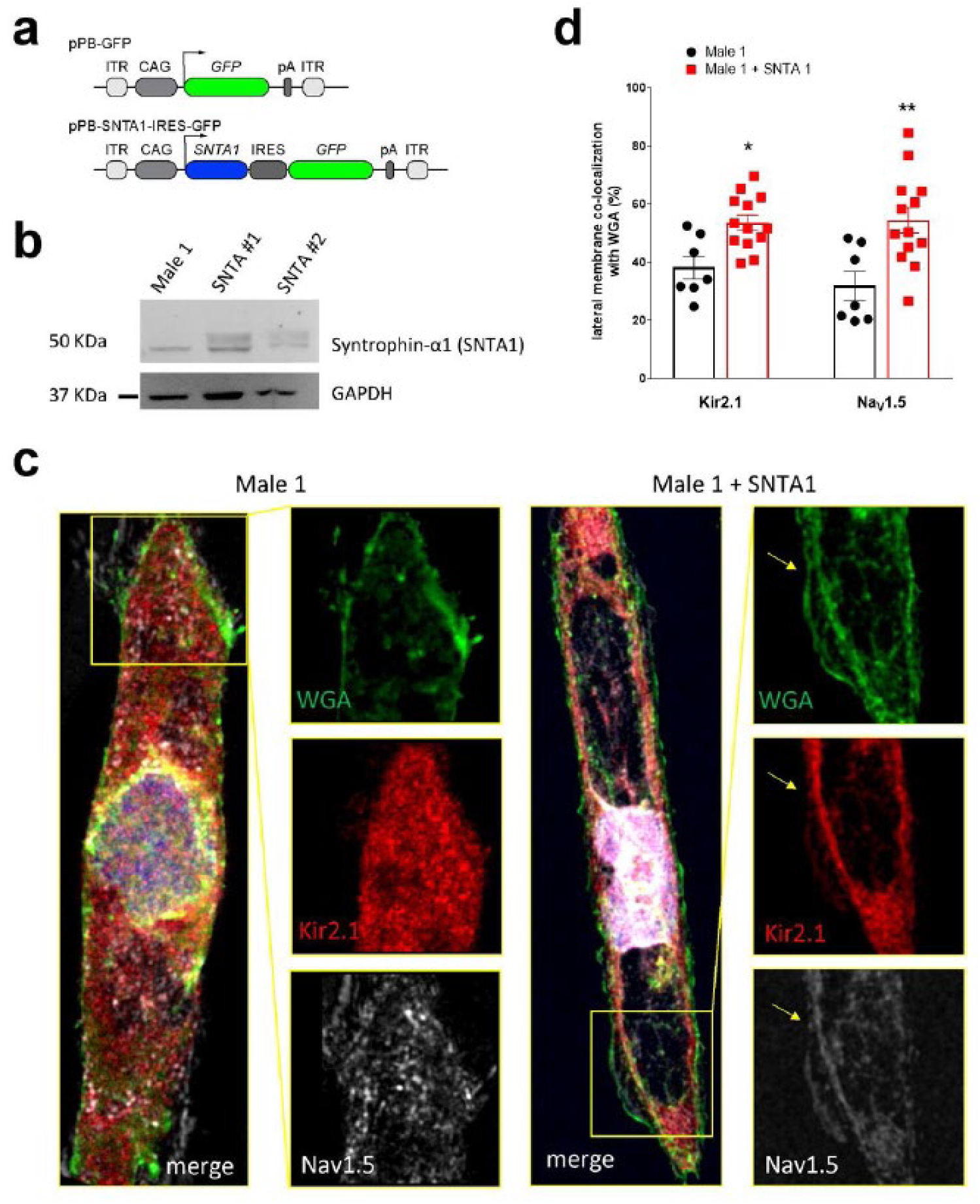
Transfection of *SNTA1* rescues membrane levels of Kir2.1 and Na_V_1.5 proteins in iPSC-CMs from Male 1 Patient. **(a)** Cartoon illustrating non-viral piggy-bac vector encoding *SNTA1* for transfection in Male 1 iPSC-CMs. *SNTA1* coding region (CDS) is driven by the CAG promoter and followed by green fluorescence protein (GFP) after an internal ribosome entry site (IRES). Control vector only expresses GFP. **(b)** Western blot for α-1-Syntrophin expression normalized with GAPDH. **(c)** Immunostaining for Kir2.1 (red), Na_V_1.5 (white) and WGA (green) in control Male 1 iPSC-CM (left) and Male 1 iPSC-CM transfected with *SNTA1*. Nuclei were stained with DAPI. Yellow arrows point to iPSC-CM membrane staining. Scale bar, 5 μm. **(d)** Quantification of Kir2.1 and Na_V_1.5 colocalization with WGA at the cell membrane shows significant increase of both Kir2.1 (*\*P* < 0.05; *n* = 7–10 cells) and Na_V_1.5 (*\*\* P* < 0.01; *n* = 7–10 cells).

### 3.7 DMD iPSC-CMs have reduced inward rectifier potassium currents

Apart from the well-described regulation of Na_V_1.5 channels by the DAPC,^10, 17^ there is evidence that this protein complex also regulates Kir2.1 inward rectifying potassium channels in *mdx* cardiomyocytes.^14^ Moreover, a pool of Na_V_1.5 channels co-localizes with Kir2.1 forming protein complexes with scaffolding proteins at the cardiomyocyte lateral membrane and intercalated disc, where they modulate each other’s surface expression.^11, 19, 20^ To test whether, in addition to *I*_Na_, the inward rectifier potassium current is also affected in iPSC-CMs from DMD patients, we compared Ba^2+^-sensitive potassium currents (*I*_K1_). In *Figure 5c* and *d*, *I*_K1_ density measured at -120 mV was significantly reduced in Male 1 (-1 ± 0.3 pA/pF) and Male 2 (-1.2 ± 0.3 pA/pF) iPSC-CMs compared to Control 1 (-3.2 ± 0.5 pA/pF). *I*_K1_ density of Control 2 cells was -2.6 ± 0.6 pA/pF (*Suppl. Figure 1i*). Changes in *I*_K1_ were highly variable in heterozygous cells, and the difference with control was not significant, likely due to the variability of expression of dystrophin (*Figure 2e*) and other proteins forming the complex.

### 3.8 Ion channel gene expression profile in male and female DMD iPSC-CMs

Previous reports have shown that when one of the DAPC components is genetically absent, other proteins of the complex are likewise down-regulated, leading to a dysfunction of the complex.^38^ To confirm whether this phenomenon occurs in both hemizygous and heterozygous DMD iPSC-CMs, we analyzed the mRNA levels, and protein expression of the cardiac ion channels Na_V_1.5 (encoded by *SCN5A* gene), Kir2.1 (encoded by *KCNJ2* gene) and Ca_V_1.2 (encoded by *CACNA1C* gene).

Consistent with what has been described for mdx mice,^10^ both hemizygous DMD iPSC-CMs showed increased *SCN5A* expression (*Suppl. Figure 7a*, *top*), also like human cardiac tissue from a Becker MD (BMD) individual (*Suppl. Figure 7a*, *bottom*). Similarly, *KCNJ2* gene expression was up-regulated in both hemizygous DMD cell lines, as well as the BMD individual (*Suppl. Figure 7b*). This suggests that the increase in cardiac *SCN5A* and *KCNJ2* mRNA levels might be a general compensatory phenomenon in DMD patients. On the other hand, consistent with the unaffected *I*_CaL_, neither *CACNA1C* nor Ca_V_1.2 were modified in either male or female DMD iPSC-CMs compared to control (*Suppl. Figure 7c*).

To test whether the decreased *I*_K1_ and *I*_Na_ in both DMD iPSC-CMs were due to reduced Na_V_1.5 and Kir2.1 protein levels, we performed Western blot experiments with total protein lysates of iPSC-CMs monolayers. In *Suppl. Figure 8a–b*, the absence of dystrophin coincided with a consistent reduction of total Na_V_1.5 protein. Surprisingly, we did not observe any change in total Kir2.1 protein. To investigate whether the reduced *I*_K1_ and *I*_Na_ in DMD iPSC-CMs was due to reduced membrane protein levels, we conducted protein biotinylation assays (*Suppl. Figure 8c–d*). Biotinylated Na_V_1.5 was significantly lower than control in the Male 2 cell line only. Biotinylated Kir2.1 was significantly reduced the hemizygous cells, consistent with the reduction in *I*_K1_. Altogether, the results presented thus far support the idea that, the absence of dystrophin in the DMD iPSC-CMs, resulted in reduced abundance of Na_V_1.5 protein in the whole cell and possibly reduced trafficking of both Na_V_1.5 and Kir2.1 to the cell membrane, as predicted from our previous work.^19–21^

The data in iPSC-CMs from the heterozygous female are more challenging. Na_V_1.5 total protein levels and biotinylated Na_V_1.5 channels were not different from control (*Suppl. Figure 8*), but the *I*_Na_ density in single iPSC-CMs was even smaller than in DMD iPSC-CMs. This, together with the lack of significance in the changes of *I*_K1_ density, total Kir2.1 protein level, and biotinylated Kir2.1, lead us to conclude that the large variability in the expression of dystrophin significantly influenced the overall results in the heterozygous cells.

### 3.9 α-1-Syntrophin expression restores electrophysiological defects in DMD iPSC-CMs

In the heart, the dystrophin-associated protein α1-syntrophin (*SNTA1*) acts as a scaffold for numerous signaling and ion channel proteins that control cardiac excitability.^33, 38, 39^ α1-syntrophin is a PDZ domain protein that co-localizes and forms a macromolecular complex (“channelosome”) with Kir2.1 and Na_V_1.5 at the sarcolemma. ^17, 19^ ^39^ ^11^ Since α1-syntrophin has been shown to modify *I*_Na_ and *I*_K1_ by enhancing membrane Na_V_1.5 and Kir2.1 membrane levels,^19^ we hypothesized that even in the absence of dystrophin, increasing α1-syntrophin should restore normal electrical function in the DMD iPSC-CMs. Therefore, we stably transfected *SNT1A* gene via piggyBac transposon-based mammalian cell expression system in Male 1 cells verifying an increase in syntrophin expression (*Figure 6a* and *b*). As illustrated in *Figure 6c*, α1-syntrophin expression increased the Kir2.1 and Na_V_1.5 protein levels in the membrane fraction as indicated by co-localization with wheat germ agglutinin (WGA) compared to controls transfected with GFP. In *Figure 7a–b*, α1-syntrophin expression resulted in a recovery of both *I*_Na_ (*Figure 8a*) and *I*_K1_ (*Figure 8b*). Consequently, as shown in *Figure 7c*, *SNT1A* transfection led to significant improvement in the electrophysiological properties of DMD iPSC-CMs. The MDP was hyperpolarized, the dV/dt_max_ and amplitude were increased and the APD_90_ was abbreviated.

**Figure 7.**
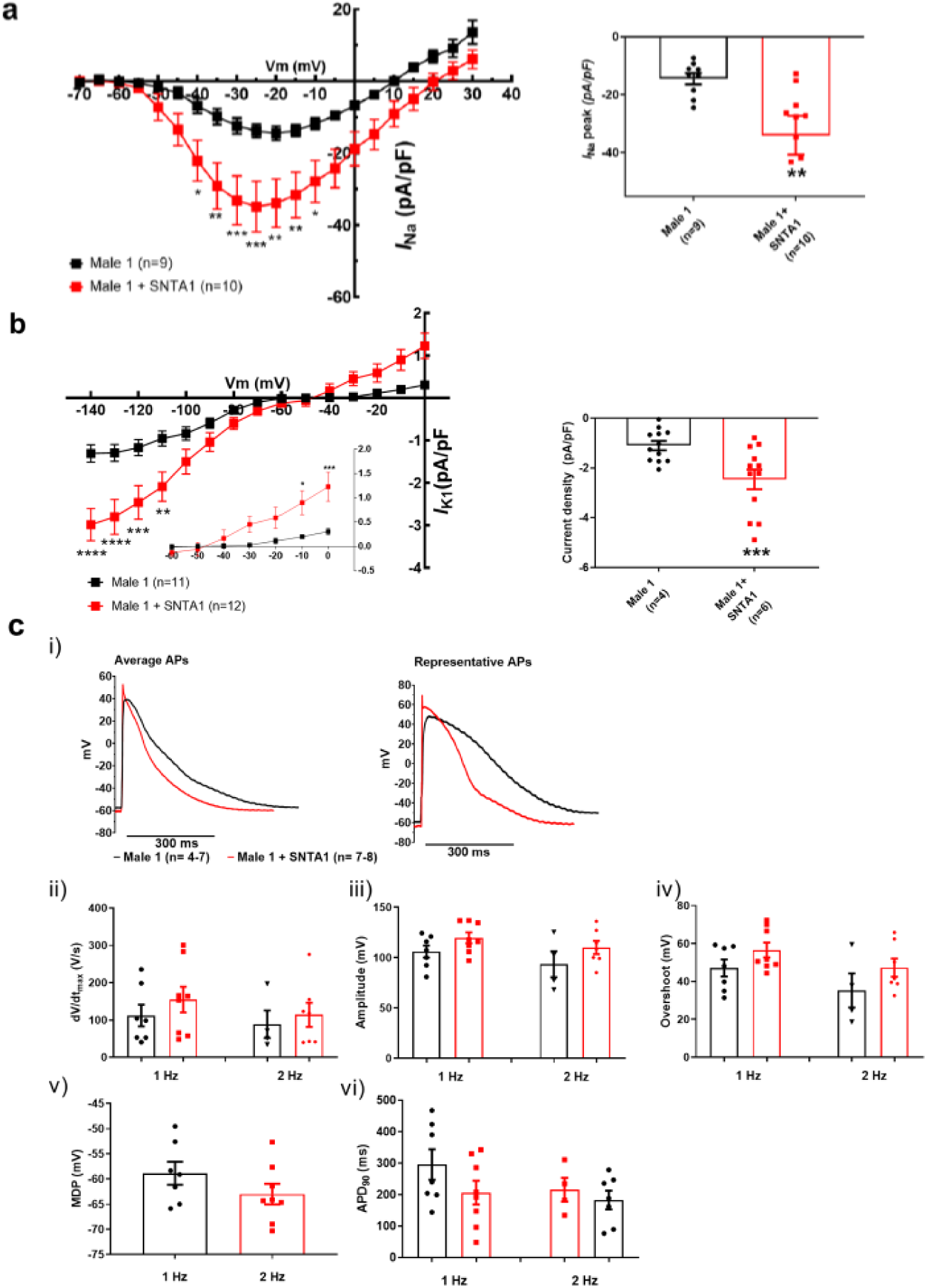
*SNTA1* expression restores the electrophysiological deficiencies in DMD iPSC-CMs. **(a–b)** Normalized current-voltage (I/V) relationships for *I*_Na_ and *I*_K1_ in Male 1 before (black) and after (red) syntrophin expression at the specified voltages. Two-way analysis of variance (ANOVA) followed by Sidak’s multiple comparisons test. Graphs show peak *I*_Na_ density at -20 mV **(a)** and peak *I*_K1_ density at -120 mV **(b)**. The inset in B highlights the increased outward component of *I*_K1_ at less negative potentials upon syntrophin expression. Two-tailed Mann-Whitney test. **(c)** Effect of syntrophin expression on AP showing: **i)** Averaged *(left)* and representative *(right)* action potential traces of ventricular-like iPSC-cardiomyocytes derived from DMD cells before (black) and after (red) syntrophin expression, **ii)** maximal AP upstroke velocity (dV/dt_max_), **iii)** Amplitude, **iv)** Overshoot, **v)** MDP and **vi)** APD_90_. Errors bars represent SEM. The *n*-values are in parentheses. \**P* < 0.05; \*\**P* < 0.01; and \*\*\**P* < 0.001.

## 4. Discussion

We demonstrate here that patient-specific iPSC-CMs recapitulated consistently the hallmark electrophysiologic features of cardiomyopathic DMD patients.^33^ In fact, mature iPSC-CMs from two hemizygous male DMD patients lacking the Dp427 isoform and a female patient heterozygous for a 5-exon deletion (Δ8–12) in the dystrophin gene have significantly reduced *I*_Na_ and *I*_K1_ densities, dV/dt_max_ and conduction velocities, as well as focal and reentrant arrhythmias. Together, these results strongly suggest that reduced excitability underlies the arrhythmogenic mechanism in DMD patients. While all patients developed severe cardiomyopathy, they also suffered frequent PVCs and ventricular tachycardia. In addition, the ECG of the heterozygous female DMD patient showed a significant left axis deviation caused by cardiac conduction defects in line with our results. In one of the male patients, ICD recordings revealed the arrhythmia deteriorating into ventricular fibrillation.^40^ Our results in patient-specific iPSC-CMs indicate that such defects are a direct consequence of a Na_V_1.5 - α1-syntrophin - Kir2.1 channelosome dysfunction produced by the disruption of the DAPC that characterizes the DMD cardiomyopathy. Remarkably, transfecting just one of the components of that complex (i.e., α1-syntrophin) in Male 1 iPSC-CMs led to channelosome recovery at the plasma membrane, with restoration of *I*_Na_ and *I*_K1_ densities, MDP, AP dV/dt_max_ and amplitude. To our knowledge, this report is first in providing a comprehensive and rigorous mechanistic demonstration of the potential causes of cardiac conduction defects and arrhythmogenesis in human DMD, substantially extending findings from animal models.^10^

ECG abnormalities can be detected in up to 60% of DMD patients,^33^ and among those, conduction defects, bradycardia, ventricular arrhythmias, and sudden death are frequent.^36^ However, despite significant progress in the understanding of the mechanisms of the skeletal muscle dystrophy, exploration of the electrophysiological consequences of the dystrophic cardiomyopathy has been slower. Until now, it has been difficult to link functional changes in individual ion channels/proteins with corresponding clinical phenotypes in inheritable ion channel diseases and cardiomyopathies such as DMD.^41^

Both Na_V_1.5 and Kir2.1 interact with the DAPC via α1-syntrophin through their respective canonical C-terminal PDZ binding domains. As shown recently, Na_V_1.5 has an additional internal PDZ-like binding domain localized at the N-terminus that also interacts with α1-syntrophin.^10, 19^ Changes in the *I*_Na_ and *I*_K1_ might alter cardiac conduction and increase the probability of premature beats like those seen in the ECG from the DMD patient.^10^ We proved here that in addition to reduced *I*_Na_, iPSC-CMs from DMD patients also have reduced *I*_K1_ and probably alterations in other proteins altogether causing pro-arrhythmic alteration in electrical impulse conduction, likely because of trafficking disruption of the α1-syntrophin-mediated macromolecular complex formed by the DACP with Kir2.1-Na_V_1.5.

Maturation of iPSC-CMs is essential for human disease modeling and preclinical drug studies.^26, 42^ Culturing iPSC-CM monolayers on soft PDMS membranes coated with Matrigel promotes cell maturation.^24^ Also, there are several reports indicating that the regulation of cell shape and substrate stiffness helps improve the contractile activity and maturation of iPSC-CMs.^25, 29^ Thus, having cells with ventricular-like action potentials and structural and electrophysiological maturity that approximates the human adult ventricular cardiomyocyte is likely to be more useful in investigating the pathophysiology of DMD patients. Therefore, here we used a micropatterning platform based on Matrigel-coated PDMS membrane^24^ for modeling single-cell cardiac electrical activity. Our findings showed that culturing single ventricular-like iPSC-CMs on micropatterned Matrigel-coated PDMS confers a cylindrical shape yielding iPSC-CMs with structural and functional phenotypes close to those in human mature cardiomyocytes.^30, 31^ Electrophysiological analyses in this scenario revealed abnormal action potential profiles in DMD iPSC-CMs, compatible with the clinical alterations observed in both male 1 and female DMD patients. The strong reduction in *I*_Na_ density yielded a significant slowing of dV/dt_max_, considered to be an indirect measure of the available functional sodium channels.^43^ Reduction in *I*_Na_ density was consistent with the relative loss of total Na_V_1.5 protein levels, and helped us explain the reduced conduction velocity in iPSC-CMs from DMD patients. Like other studies,^17, 44^ we did not find any change in Cx43 protein levels.

QRS widening and QTc prolongation displayed on the ECGs from the DMD patients are likely related to the changes in functional expression of Na_V_1.5 and Kir2.1 we have observed in their iPSC-CMs. Both QRS widening and QT dispersion are risk factors for arrhythmias in patients with DMD, and have been implicated in the genesis of ventricular arrhythmias.^45^ Interestingly some of the AP parameters of the hemizygous Male 2 iPSC-CMs, including dV/dtmax, AP amplitude and overshoot (Table 1), were substantially more reduced than Male 1 and the heterozygous female iPSC-CMs. Such differences are possibly due to the specific mutation in the dystrophin gene. Thus, depending on the mutation in the dystrophin gene each male or female DMD patient might develop different types or levels of cardiac electrical dysfunction and life-threatening arrhythmias.

*I*_Na_ reduction coincided with *I*_K1_ reduction in both hemizygous DMD iPSC-CMs, supporting the idea that both channels require PDZ-mediated interaction with components of the DAPC to modulate reciprocally their proper expression._10, 18_ It is likely that the reduced *I*_K1_ in the DMD iPSC-CMs contributed to the reduced dV/dt_max_, although the MDP in the iPSC-CMs from the two dystrophic patients was like control. In this regard, it is important to note that the relationship between MDP and *I*_Na_ availability is highly nonlinear in such a way that a very small reduction in MDP is expected to result in substantial reduction in sodium current during the action potential upstroke.^46^ Regardless, the biotinylation experiments demonstrated that Kir2.1 levels at the membrane were significantly lower in both DMD iPSC-CMs with respect to the control. The elevated *SCN5A* and *KCNJ2* mRNA levels excluded the possibility that a decrease in gene expression was responsible for the protein loss, and therefore, to smaller *I*_Na_ and *I*_K1_ densities in the DMD iPSC-CMs. This somehow contrasts with reports in *mdx*^5cv^ mouse hearts, where the Na_V_1.5 mRNA levels remained unchanged with a strong reduction in the Na_V_1.5 protein levels.^10^ As such, the reduction in the Na_V_1.5 and Kir2.1 protein levels could be related to ubiquitylation and proteasome degradation as suggested previously in studies in dystrophin-deficient *mdx*^5cv^ mice.^47^ However, our results in DMD iPSC-CMs strongly suggest that disruption of the DAPC due to lack of dystrophin significantly impairs ion channel expression and function.^10, 15, 16^ Specifically, we demonstrate that the decrease in ion channel current densities is the result of Na_V_1.5 and Kir2.1 trafficking and membrane targeting defects directly derived from the absence of dystrophin. Such defects can be completely reverted by α1-syntrophin expression, as demonstrated by increases in *I*_Na_ and *I*_K1_, and restoration of MDP, action potential upstroke velocity and action potential amplitude, as well as APD abbreviation. On the other hand, the fact that both *I*_Na_ and *I*_K1_ are only partially reduced in the DMD iPSC-CMs suggests the presence of different pools of Na_V_1.5 and Kir2.1 channels that do not depend on DAPC integrity. Altogether, our results support the idea that DMD cardiomyopathy results in ion channel dysfunction that predisposes the dystrophic ventricular myocardium to arrhythmia with potentially lethal consequences.

Previous reports indicate that although heterozygous DMD females, have negligible skeletal muscle symptoms, they are not free of cardiac involvement.^48^ For example, the clinical expression of the X-linked DMD cardiomyopathy of heterozygous females increases with age.^48^ The female patient represented in this study suffered from a relative severe phenotype, characterized by skeletal myopathy and cardiomyopathy, which could be explained by a malignant mutation disrupting the N-terminal of the dystrophin gene. One could assume that one gene of dystrophin should produce enough dystrophin to preserve function in multinucleated skeletal muscle of females._49_ Unexpectedly, we found that *I*_Na_ density in iPSC-CMs from the heterozygous female was even more reduced compared to hemizygous iPSC-CMs. Interestingly, the QRS duration was significantly prolonged on the ECG from the heterozygous female compared to the hemizygous patient (see Figure 1), suggestive of a more dramatic loss-of-function effect on Na_V_1.5 in heterozygous females. Probably this is related to the heterogeneity seen in immunostaining studies where some heterozygous female cells express normal dystrophin levels while others show absence or very low expression likely due to random X-inactivation of the WT allele.^23^ Because of random inactivation of one of the X chromosomes, heterozygous females should constitute a mosaic of 2 or more cell types dramatically differing in the extent of dystrophin expression. Thus, it would not be surprising that females with DMD are more prone to suffer arrhythmias because of spatial electrical inhomogeneity due to variable expression of the mutant allele. The heterogeneity in dystrophin expression has been also observed in canine *carrier* models of X-linked dystrophy, which exhibit a cardiac mosaic pattern, where dystrophin in each myocyte is either fully expressed or absent.^50^ Nevertheless, the importance of abnormal cardiac measures in heterozygous females who harbor mutations in the dystrophin gene remains debatable.^51^

Even though *I*_Na_ density was substantially reduced in the heterozygous iPSC-CMs, neither the total Na_V_1.5 protein levels nor the biotinylated Na_V_1.5 showed any changes. Probably, the variable expression of dystrophin in female individuals results in variable Na_V_1.5 protein levels, while Kir2.1 expression and function are modulated positively to help trafficking of the few pools of Na_V_1.5 channels belonging to the remaining DAPC. Another possibility that might explain the reduced *I*_Na_ in heterozygous iPSC-CMs is that the cells may lack a suitable compensatory response due to DAPC disorganization and malfunction. The chimeric nature of the dystrophin mutation in those cells likely makes it more difficult to support a compensatory mechanism than the complete absence of the DAPC complex as it occurs in dystrophic cells. Nonetheless, the very reduced *I*_Na_ and slowed CV reported in the present study perfectly correlates with the clinical data from the heterozygous female patient. Prolonged QRS duration is evidence of slowed ventricular activation and inhomogeneous conduction and might be associated with rotor activity as observed in the *female* iPSC-CMs monolayers, which is considered a substrate for reentrant ventricular tachycardia.^52^ This becomes important because although controversial, heterozygous females may have an age-related increased risk of cardiac conduction disease and sudden death; in female patients of X-linked Emery-Dreifuss muscular dystrophy cardiac alterations typically occur late in life.^53^

## 5. Limitations

We have derived data from experiments conducted in iPSC-CMs from patients who carry independent dystrophin mutations and two unrelated controls, which may be a potential limitation of our study. The original study design included siblings for each DMD cell line. However, getting more experimental groups from the same family was not possible. Nevertheless, both DMD lines lack dystrophin, which gives credence to the idea that loss of dystrophin is important to the shared electrophysiological phenotype independently of the specific mutation. Further, we show new insight into how heterozygous DMD females might show a wide range of cardiac involvement, ranging from asymptomatic to severely impaired electrical cardiac function, particularly the highly reduced *I*_Na_ leading to slowing of conduction velocity, which is reflected on the ECG from the female patient. Thus, together with the structural alterations, the electrophysiological changes may contribute to left ventricular dysfunction in female DMD patients.^54^ However, the impact of the finding that the female carrier of the mutation presents a decrease in *I*_Na_ is somehow mitigated by the fact that since she carries a different mutation, it is difficult to define how the reduction of the *I*_Na_ in the female carrier compares with the reduction observed in the affected individuals.

iPSC-CMs still show significant differences with adult ventricular cardiomyocytes and are still far from recapitulating chamber-specific and layer specific electrical phenotypes of the normal or dystrophic heart. In addition, we cannot generalize our results to patients with different dystrophic gene mutations, such as those underlying Becker muscular dystrophy, which lead to partially truncated dystrophins and may retain specific functional properties of full-length dystrophin. However, enrolling a Becker MD patient was not possible. Also, our syntrophin-mediated rescue experiments were limited to the Male 1 iPSC-CMs line. While caution should be exerted when attempting to extrapolate to the other two DMD cell lines, it is important to note that the functional defects in the Na_V_1.5-Kir2.1 channelosome were very similar in the iPSC-CMs from all three patients, which gives credence to our interpretation.

## Data Availability

Authors will make materials, data and associated protocols promptly available to readers without undue qualifications upon request.

### Funding

This work was supported by National Institutes of Health R01 HL122352 grant; “la Caixa” Banking Foundation (HR18-00304); Fundación La Marató TV3: *Ayudas a la investigación en enfermedades raras* 2020 (LA MARATO-2020); and Instituto de Salud Carlos III to JJ. American Heart Association postdoctoral fellowship 19POST34380706s to J.V.E.N. Israel Science Foundation to OB and MA [824/19]. Rappaport grant [01012020RI]; and Niedersachsen Foundation [ZN3452] to OB; and US-Israel Binational Science Foundation (BSF) to OB and TH [2019039]. National Institutes of Health R01 AR068428 to DM and US-Israel Binational Science Foundation Grant [2013032] to DM and OB.

## Author Contributions

E.N.J.V. and J.J. designed research, discussed results and strategy. E.N.J.V., conducted the experiments to characterize the DMD iPSC-CMs; M.A. cared for the DMD patients and provided the skin biopsies; A.M. conducted the patch-clamp experiments showing *SNTA1* rescue of electrical properties of DMD iPSC-CMs; M.L.V.P. differentiated, transfected and sorted the iPSC-CMs used in *SNTA1* rescue and helped in immunolocalization experiments; F.M.C.U. designed and generated the *SNTA1* piggyBac transposon-based constructs and conducted immunolocalization experiments; O.B. reprogrammed and characterized the iPSC lines. A.J.C. provided technical assistance in the generation of all the iPSC-CMs lines. A.J.C., G.G.S., A.M.D.R., and D.P.B. conducted, collected and analyzed the experiments. E.N.J.V. and J.J. wrote the manuscript. Critical revision of the manuscript: all authors.

## Acknowledgments

We thank Dr. Adam Helms for help in the implementation of the micropattern technology. We also thank Joseph R. Dickens from the CSCAR-University of Michigan for his valuable guidance in the statistical analysis. Dov Freimark MD (from the Leviev Heart Center, Sheba Medical Center, Tel Hashomer, and Tel Aviv University, Israel) supported this study by caring for and enrolling the female patient. We thank Dr. Giovanna Giovinazzo and the staff of the CNIC Pluripotent Cell Unit for their help in processing the iPSCs used in the syntrophin-mediated rescue experiments. We are grateful to patients who despite having a lethal disease agreed to undergo skin biopsy for the sake of science.

## Conflict of interest

<u>JJ</u>. Elsevier royalties; Stembiosys Scientific Advisory Board, stock options; <u>AMdR</u>: Stembiosys consulting fees and stock options; TJH, Stembiosys Scientific Advisory Board, consulting fees and stock options; All other authors have no conflicts to declare.

## SUPPLEMENTARY DATA

### Supplementary Figures

**Supplemental Fig. 1.**
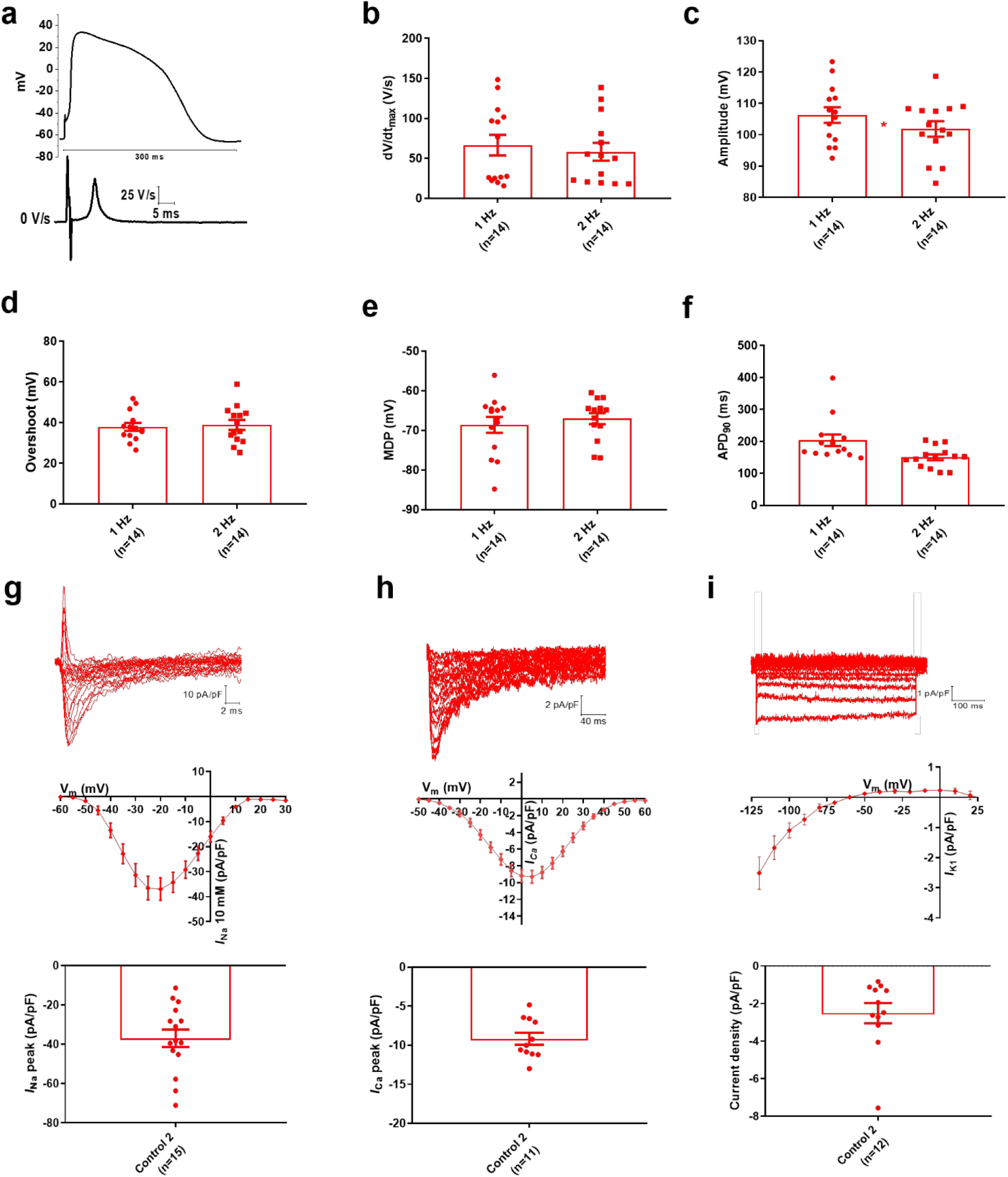
Electrophysiological analysis in Control 2 (human foreskin-derived BJ iPSC-CMs). (**a**) Representative AP trace of ventricular-like control BJ iPSC-CMs obtained at 1 Hz of pacing. *Inset*. First derivative with respect to time (dV/dt). (**b–f)** Action potential properties. Recordings at 1 and 2 Hz were similar to those obtained from the healthy donor patient derived-iPSC-CMs (Control 1). (**g-i)** Current traces, I/V curves, and normalized current densities for NaV1.5, CaV1.2, and Kir2.1 ion channels, respectively. Data obtained from the control BJ iPSC-CMs (Control 2) were similar to the other control iPSC-CMs.

**Supplemental Fig. 2.**
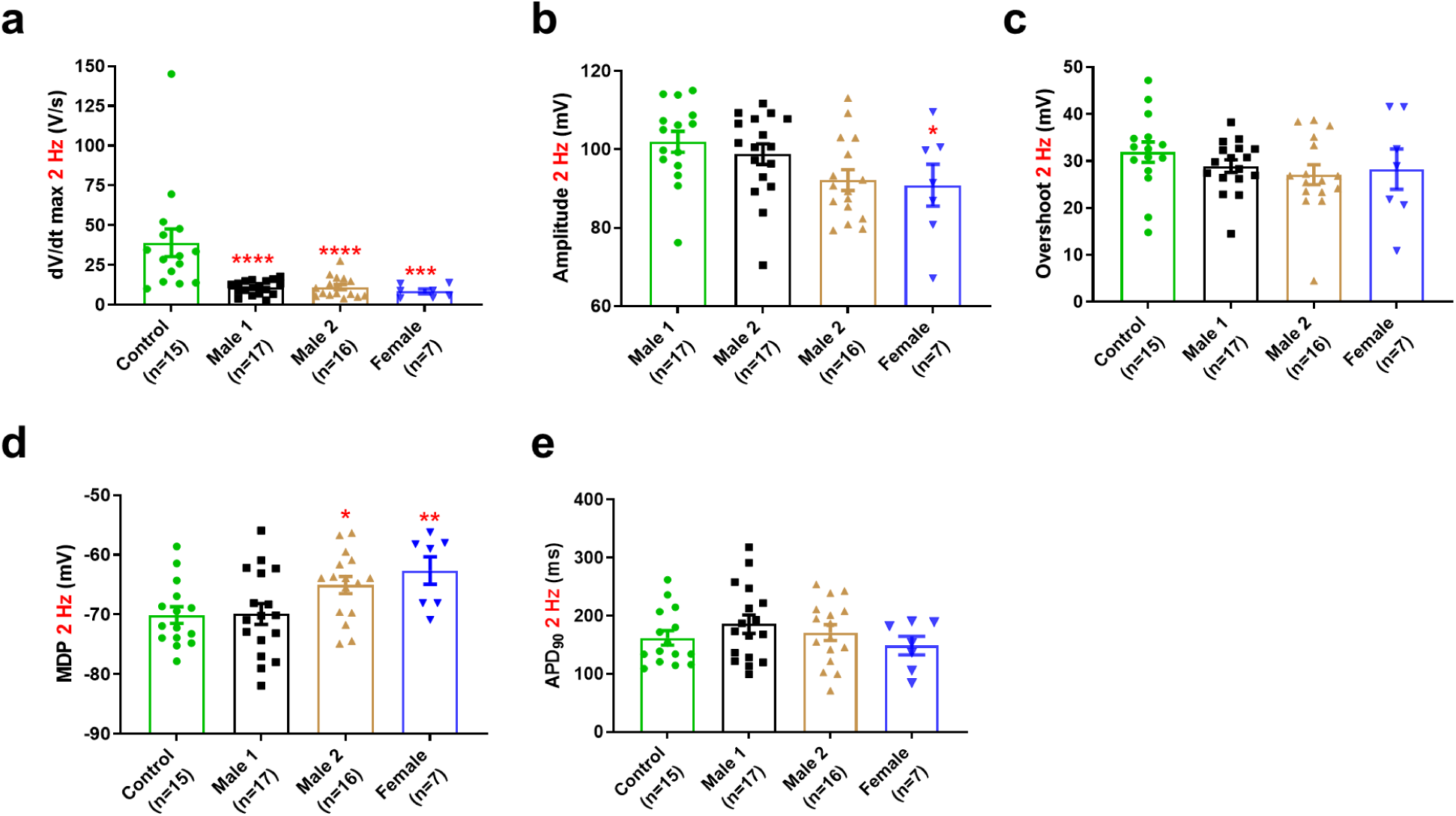
Action potential properties of control, hemizygous and heterozygous DMD iPSC-CMs paced at 2 Hz. **(a)** Maximal AP upstroke velocity was reduced in both hemizygous (\*\*\*\**P* < 0.0001), and the heterozygous female (\*\*\**P* = 0.0002) iPSC-CMs compared to control. **(b–d)** Overshoot values were similar among all tested groups (*P* = 0.2413 for Male 1 cells, *P* = 0.1121 for Male 2 cells, and *P* = 0.4115 for *female* cells). Amplitude and MDP were statistically significant affected in the Male 2 iPSC-CMs (\**P* = 0.0109 and \**P* = 0.0267, respectively) compared to control, while the heterozygous iPSC-CMs showed a more depolarized RMP (\*\**P* = 0.0081). **(e)** APD90 was similar in all tested iPSC-CMs (*P* = 0.3699, *P* = 0.5196, and *P* = 0.8366 for Male 1, Male 2, and heterozygous iPSC-CMs, respectively). Cells plated on micropatterns were paced at 2 Hz. Two-tailed Mann-Whitney test. Errors bars represent s.e.m. The *n*-values are in parentheses.

**Supplemental Fig. 3.**
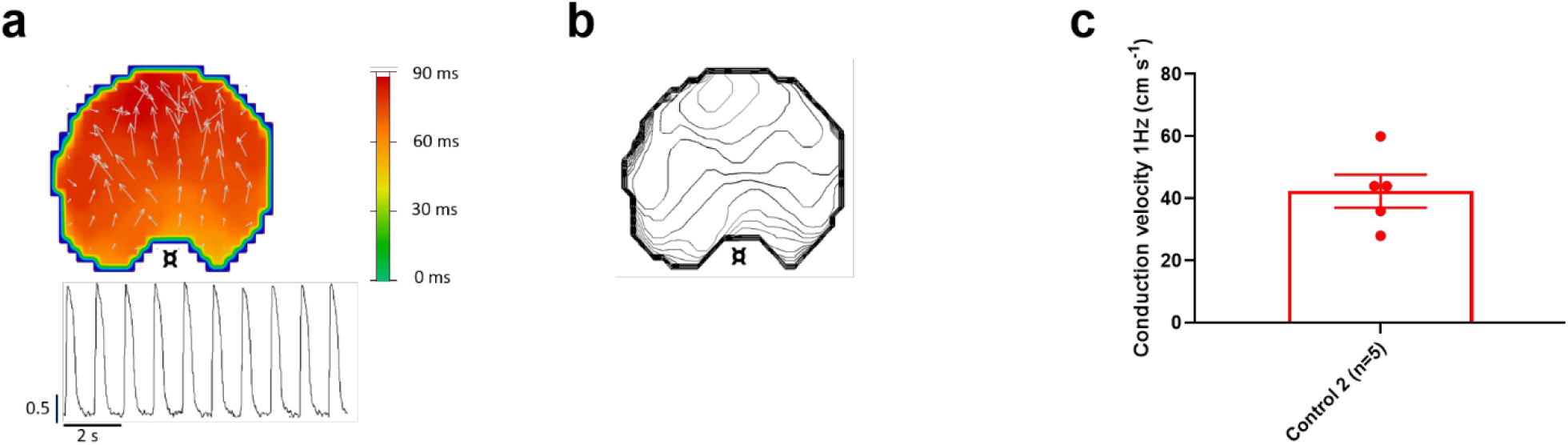
Conduction velocity in control BJ iPSC-CM (Control 2) monolayer. **(a)** Activation maps of action potential propagation at 1 Hz. Each color represents a different activation time with time zero appearing in green (**¤** indicates the location of the stimuli for each monolayer). White vectors (↑) are a measure of local velocity and direction of the wave. *Inset.* Representative optical APs evoked by external stimulation at 1 Hz. **(b)** Averaged 2-s contour isochrone maps for the representative monolayer in A. **(c)** Bar graph of conduction velocity in the additional control BJ monolayers. Errors bars represent SEM. The *n*-values are in parentheses.

**Supplemental Fig. 4.**
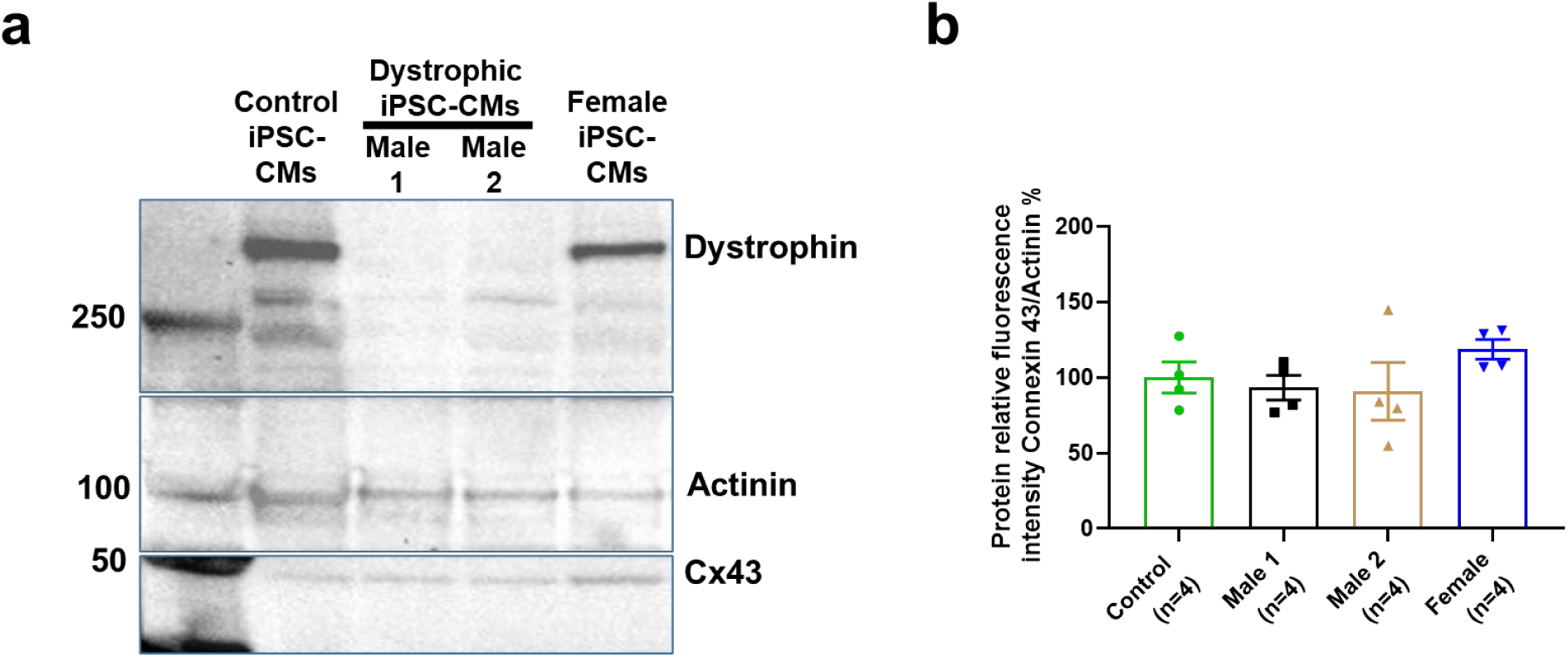
Cx43 expression level in control, hemizygous and heterozygous DMD iPSC-CMs. **(a)** Typical Western blot for connexin43 expression. About 50k cells were collected to quantify total dystrophin, connexin43 and actinin levels in iPSC-CMs. **(b)** Scatter plots of Cx43 detected in control, hemizygous and heterozygous DMD iPSC-CMs. Cx43 protein levels normalized to actinin (loading control) resulted to be similar in all tested groups (*P* = 0.8857 for Male 1, *P* = 0.6857 for Male 2, and *P* = 0.1143 for female iPSC-CMs). Two-tailed Mann-Whitney test. Errors bars represent s.e.m. The *n*-values are indicated in parentheses after the name of each group.

**Supplemental Fig. 5.**
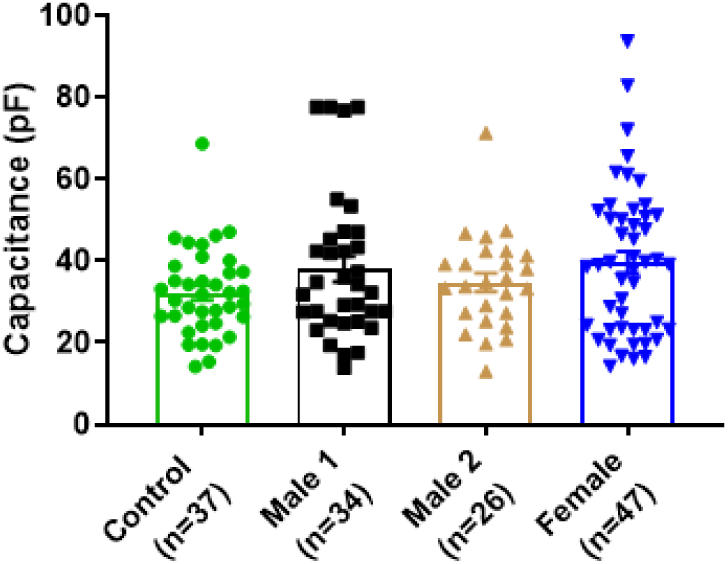
Cell capacitance. No statistically significant differences in size were observed among control (32 ± 2 pF), female (39 ± 3 pF; *P* = 0.0715) and DMD iPSC-CMs (38 ± 3 pF, *P* = 0.3257 for Male 1; and 35 ± 2 pF, *P* = 0.3703 for Male 2 iPSC-CMs). Two-tailed Mann-Whitney test. Errors bars represent s.e.m. The *n*-values are indicated in parentheses after the name of each group.

**Supplemental Fig. 6.**
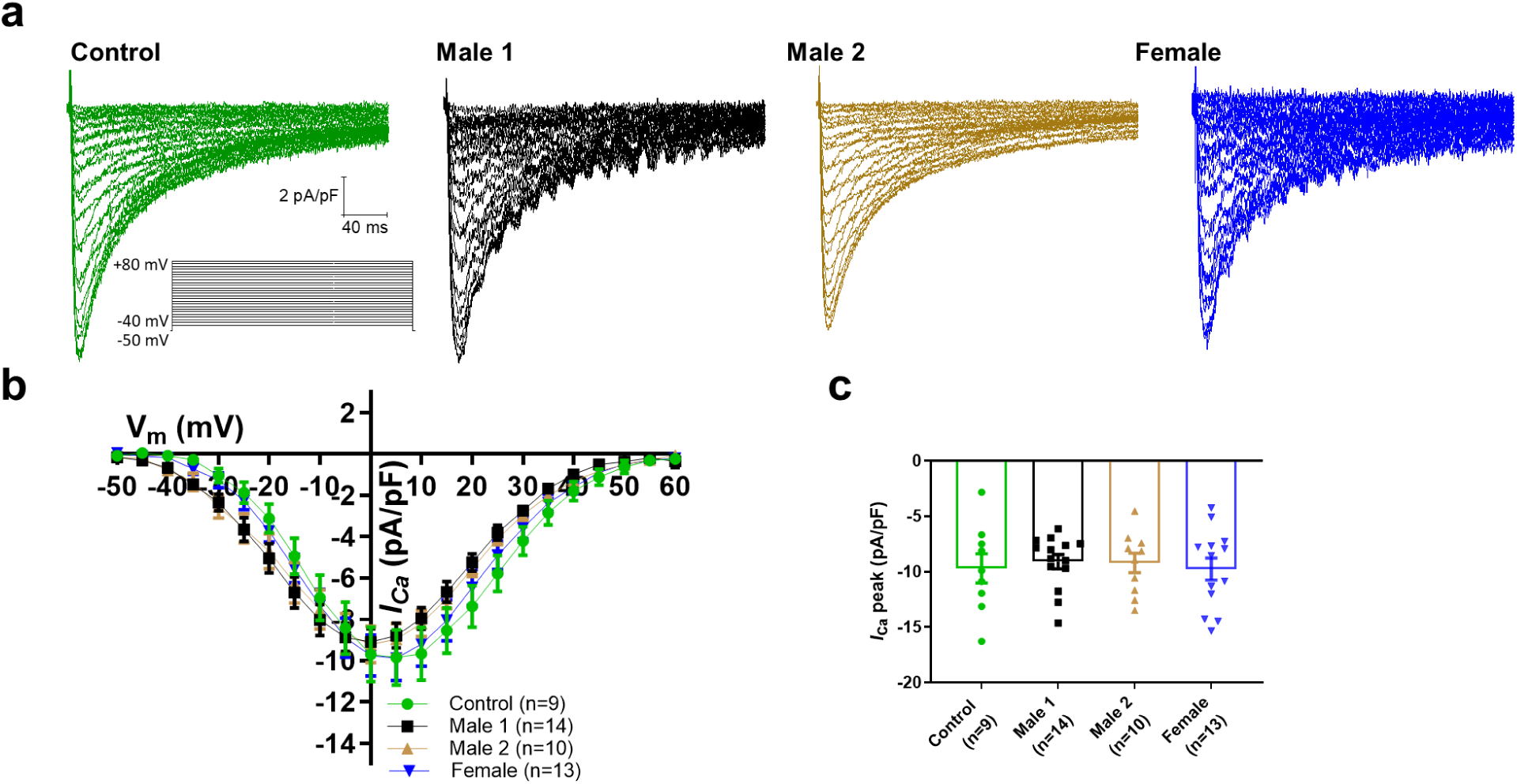
Calcium channel properties in control, DMD and female iPSC-CMs. **(a)** Original calcium current traces obtained from all iPSC-CMs elicited by depolarizing potential as shown in the *inset*. (**b**) Current-voltage relationships showing no significant differences among all tested groups. Two-way ANOVA followed by Sidak’s multiple comparisons. **(c)** Comparison of normalized current densities from all iPSC-CMs groups. The current values at 0 mV were similar in all analyzed cells (*P* = 0.5571 for Male 1, *P* = 0.8421 for Male 2, and *P* > 0.9999 for female iPSC-CMs). Two-tailed Mann-Whitney test. Errors bars represent s.e.m. The *n*-values are indicated in parentheses after the name of each group.

**Supplemental Fig. 7.**
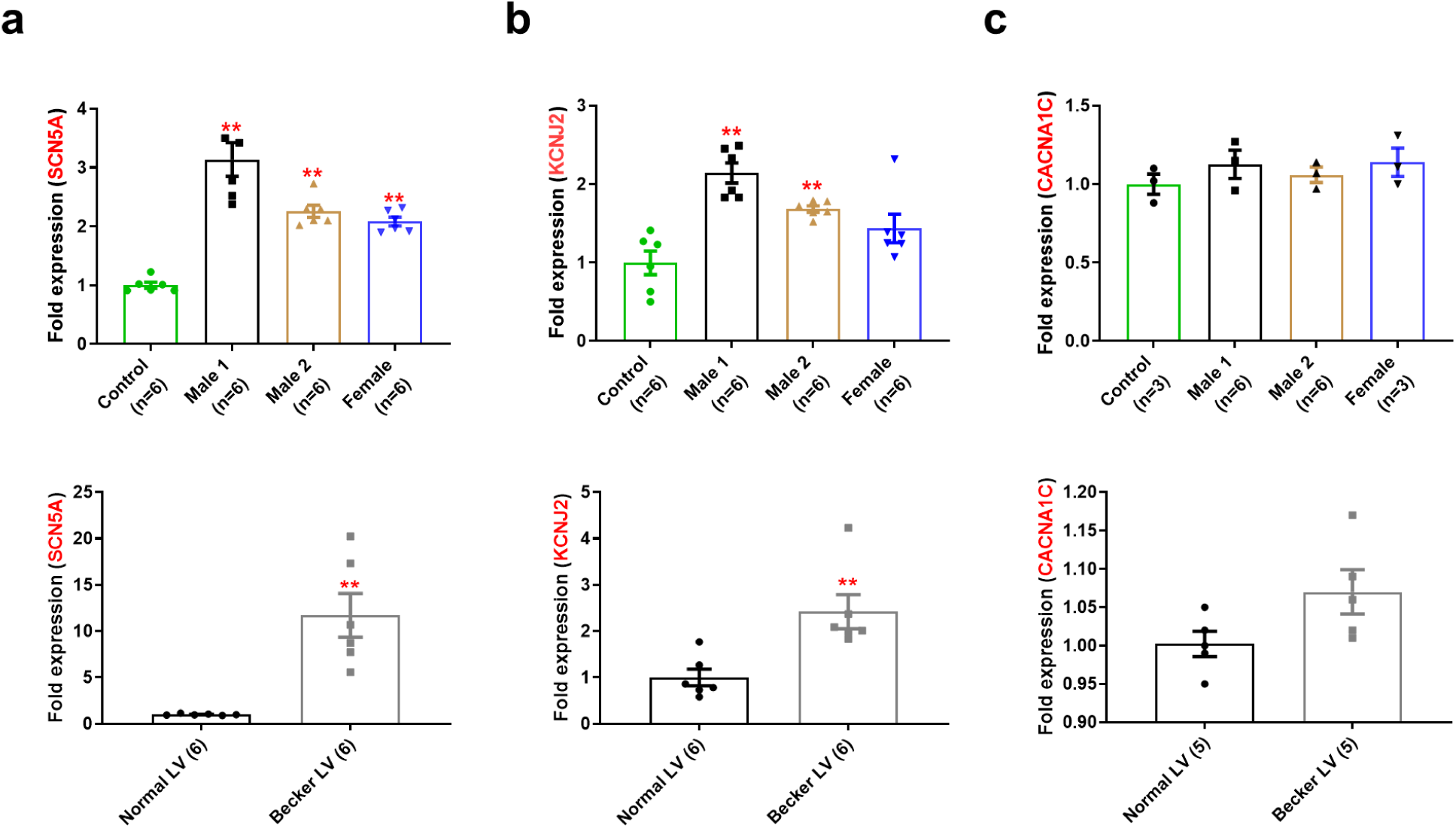
*SCN5A*, *KCNJ2* and *CACNA1C* mRNA expression in control, DMD, and female iPSC-CMs. **(a)** *SCN5A* mRNA expression was increased in iPSC-CMs from hemizygous and heterozygous DMD individuals (top), as well as in the human left ventricle heart tissue from a Becker MD individual compared to a healthy subject (bottom). **(b)** *KCNJ2* mRNA levels were higher in both hemizygous and heterozygous iPSC-CMs (top), like those found in human left ventricle heart tissue from Becker DM patients (bottom) when compared to the corresponding control. **(c)** *CACNA1C* mRNA expression was not significant different among tested groups from either iPSC-CMs (top) or left ventricle tissues (bottom). mRNA levels were determined by qRT-PCR and calculated by the comparative Ct method (2^-ddCt^) normalized to the internal control 18s rRNA. Errors bars represent SEM. The *n*-values are in parentheses. Two-tailed Mann-Whitney test. \*\**P* < 0.005.

**Supplemental Figure 8.**
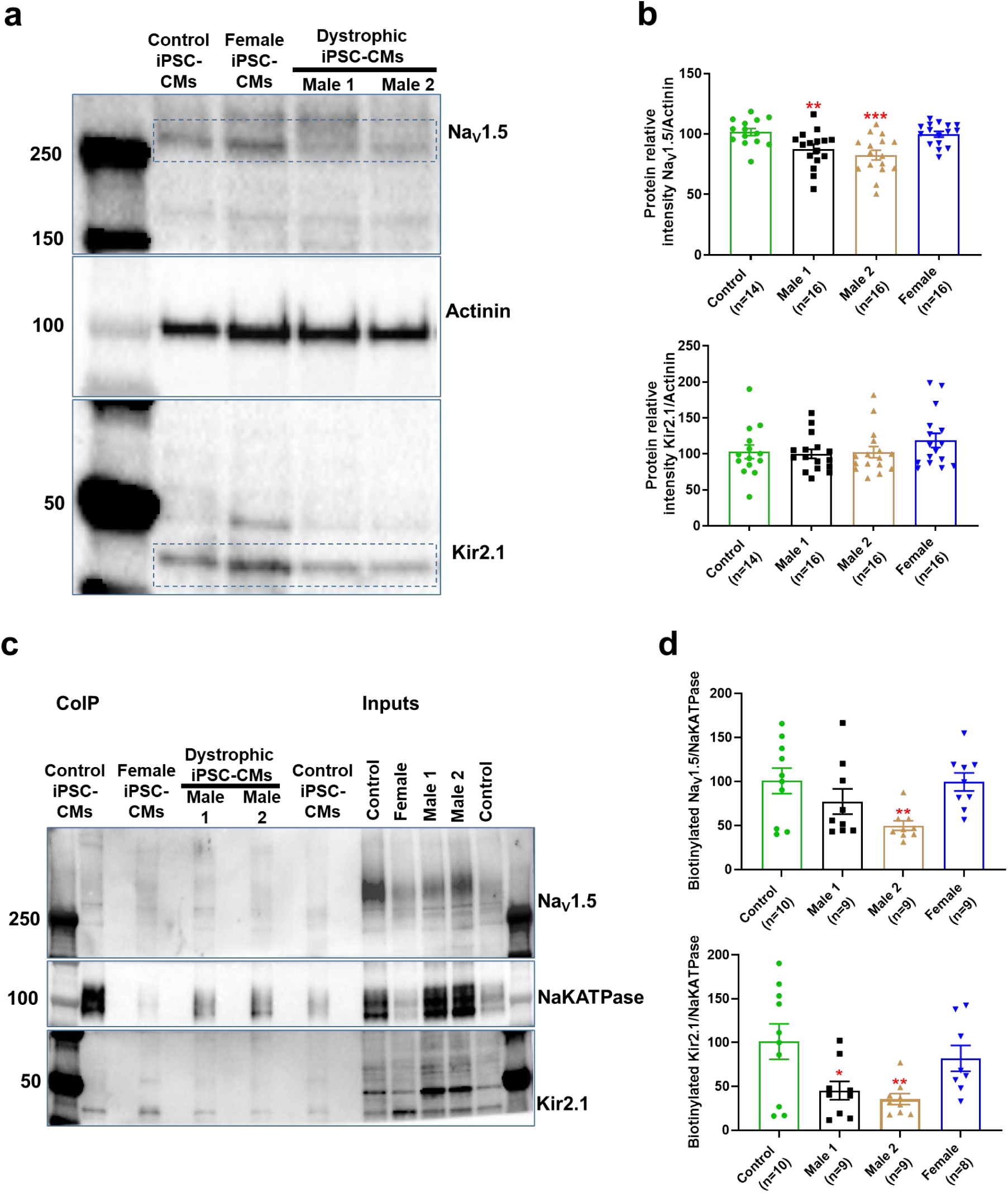
NaV1.5 protein level is significantly reduced in patient-specific DMD iPSC-CMs. **(a)** Representative Western blot for each antibody used. The bands within the blue rectangles at ∼250 KDa and below 50 KDa correspond to NaV1.5 and Kir2.1, respectively. About 50K cells were collected to quantify total NaV1.5, Kir2.1 and actinin levels in control, heterozygous and hemizygous DMD cells. **(b)** Scatter plots of NaV1.5 and Kir2.1 detected in control, *female* and DMD iPSC-CMs. NaV1.5 and Kir2.1 protein levels were normalized to actinin (loading control). **(c)** Representative Western blot after biotinylation and protein precipitation with streptavidin magnetic beads. **(d)** Scatter plots of biotinylated NaV1.5 and Kir2.1 from control, *female*, and DMD iPSC-CMs. Fifty μg of biotinylated protein was loaded. Errors bars represent SEM. The *n*-values are in parentheses. Two-tailed Mann-Whitney test. \*\*\**P* < 0.001, \*\**P* < 0.01, and \**P* < 0.05

### Supplementary Tables

**Supplemental Table 1.** Action potential parameters of iPSC-CMs paced at 1 and 2 Hz.

| Group | dV/dt<sub>max</sub> | Overshoot | Amplitude | MDP | APD<sub>90</sub> | n |
| --- | --- | --- | --- | --- | --- | --- |
| **1 Hz** |  |  |  |  |  |  |
| Control 1 | 32 ± 5 | 31 ± 2 | 101 ± 3 | -70 ± 2 | 171 ± 17 | 13 |
| Male 2 | 8 ± 1* | 23 ± 2* | 85 ± 2* | -64 ± 2 | 171 ± 19 | 12 |
| Male 1 | 11 ± 1* | 31 ± 1 | 103 ± 2 | -70 ± 2 | 218 ± 21 | 15 |
| Female | 12 ± 2 | 28 ± 3 | 92 ± 4 | -63 ± 2* | 169 ± 22 | 9 |
| **2 Hz** |  |  |  |  |  |  |
| Control 1 | 39 ± 9 | 32 ± 2 | 102 ± 3 | -70 ± 1 | 162 ± 12 | 15 |
| Male 2 | 11 ± 2**** | 27 ± 2 | 92 ± 3* | -65 ± 1* | 171 ± 14 | 16 |
| Male 1 | 11 ± 1**** | 29 ± 1 | 99 ± 3 | -70 ± 2 | 186 ± 16 | 17 |
| Female | 9 ± 1*** | 28 ± 4 | 91 ± 5 | -63 ± 2** | 149 ± 16 | 7 |

One-way ANOVA followed by Dunnett's multiple comparisons test. Values are expressed as mean ± SEM.
\*\*\*\* $P < 0.0001$ , \*\*\* $P = 0.0002$ , \*\* $P = 0.0081$ , and \* $P < 0.05$

**Supplemental Table 2.** Action potential parameters of iPSC-CMs paced at 1 and 2 Hz.

| Group | dV/dt<sub>max</sub> | Overshoot | Amplitude | MDP | APD<sub>90</sub> | n |
| --- | --- | --- | --- | --- | --- | --- |
| **1 Hz** |  |  |  |  |  |  |
| Control 2 | 66 ± 12 | 38 ± 2 | 106 ± 2 | -69 ± 2 | 204 ± 18 | 14 |
| Male 2 | 9.6 ± 2*** | 23 ± 2**** | 88 ± 3*** | -64 ± 2 | 171 ± 19 | 12 |
| Male 1 | 11 ± 1**** | 32 ± 1 | 103 ± 2 | -70 ± 2 | 218 ± 21 | 15 |
| Female | 12 ± 2*** | 28 ± 3* | 92 ± 4** | -63 ± 2 | 169 ± 22 | 9 |
| **2 Hz** |  |  |  |  |  |  |
| Control 2 | 58 ± 11 | 39 ± 2 | 102 ± 2 | -67 ± 1 | 150 ± 9 | 14 |
| Male 2 | 11 ± 2**** | 27 ± 2** | 92 ± 3 | -65 ± 1 | 171 ± 14 | 16 |
| Male 1 | 11 ± 1**** | 29 ± 1** | 99 ± 3 | -70 ± 2 | 186 ± 16 | 17 |
| Female | 9 ± 1*** | 28 ± 4* | 91 ± 5 | -63 ± 2 | 149 ± 16 | 7 |

One-way ANOVA followed by Dunnett's multiple comparisons test. Values are expressed as mean ± SEM.
\*\*\*\* $P < 0.0001$ , \*\*\* $P < 0.0007$ , \*\* $P < 0.0089$ , and \* $P < 0.05$ .

**Supplemental Table 3.** Action potential parameters of iPSC-CMs at 1 and 2 Hz, Control 1 vs Control 2.

| Group | dV/dt<sub>max</sub> | Overshoot | Amplitude | MDP | APD<sub>90</sub> | *n* |
| --- | --- | --- | --- | --- | --- | --- |
| **1 Hz** |  |  |  |  |  |  |
| Control 1 | 44 ± 13 | 31 ± 2 | 101 ± 3 | -70 ± 2 | 171 ± 17 | 13 |
| Control 2 | 66 ± 12 | 38 ± 2 | 106 ± 2 | -69 ± 2 | 204 ± 18 | 14 |
| **2 Hz** |  |  |  |  |  |  |
| Control 1 | 39 ± 9 | 32 ± 2 | 102 ± 3 | -70 ± 1 | 162 ± 12 | 15 |
| Control 2 | 58 ± 11 | 39 ± 2 | 102 ± 2 | -67 ± 1 | 150 ± 9 | 14 |

One-way ANOVA followed by Dunnett's multiple comparisons test. Values are expressed as mean ± SEM.

**Supplemental Table 4.** Biophysical parameters of DMD, and female iPSC-CMs vs Control 1.

| **Activation** |  |  |  |  |  |
| --- | --- | --- | --- | --- | --- |
|  | V<sub>50</sub> | *k* | V<sub>rev</sub> | Peak current density | *n* |
| Na<sup>+</sup> currents | mV | mV | mV | pA/pF |  |
| Control 1 | -29 ± 1 | 3 ± 1 | 17 ± 1 | -27 ± 3 | 14 |
| Male 1 | -31 ± 1 | 4 ± 1 | 16 ± 1 | -14 ± 1**** | 11 |
| Male 2 | -28 ± 1 | 3 ± 1 | 17 ± 2 | -15 ± 1**** | 9 |
| Female | -27 ± 1 | 4 ± 1 | 14 ± 1 | -11 ± 1**** | 22 |
| **Ca<sup>2+</sup> currents** |  |  |  |  |  |
| Control 1 | - 8 ± 1 | 8 ± 1 | 54 ± 3 | -10 ± 1 | 9 |
| Male 1 | -16 ± 1 | 8 ± 1 | 54 ± 2 | - 9 ± 1 | 14 |
| Male 2 | -14 ± 1 | 9 ± 1 | 57 ± 1 | - 9 ± 1 | 10 |
| Female | -12 ± 1 | 7 ± 1 | 54 ± 2 | -10 ± 1 | 13 |

**Supplemental Table 5.** Biophysical parameters of DMD, and female iPSC-CMs vs Control 2.

| Activation |  |  |  |  |  |
| --- | --- | --- | --- | --- | --- |
| | $V_{50}$ | $k$ | $V_{rev}$ | Peak current density | $n$ |
| Na<sup>+</sup> currents | mV | mV | mV | pA/pF |  |
| Control 2 | -34 ± 1 | 4 ± 1 | 15 ± 1 | -38 ± 1 | 15 |
| Male 1 | -31 ± 1 | 4 ± 1 | 16 ± 1 | -13 ± 1**** | 11 |
| Male 2 | -28 ± 1 | 3 ± 1 | 17 ± 2 | -15 ± 1**** | 9 |
| Female | -27 ± 1 | 4 ± 1 | 14 ± 1 | -11 ± 1**** | 22 |
| Ca<sup>2+</sup> currents |  |  |  |  |  |
| Control 2 | -11 ± 1 | 8 ± 1 | 55 ± 1 | -10 ± 1 | 11 |
| Male 1 | -16 ± 1 | 8 ± 1 | 54 ± 2 | -9 ± 1 | 14 |
| Male 2 | -14 ± 1 | 9 ± 1 | 57 ± 1 | -9 ± 1 | 10 |
| Female | -12 ± 1 | 7 ± 1 | 54 ± 2 | -10 ± 1 | 13 |

**Supplemental Table 6.** Biophysical parameters of iPSC-CMs, Control 1 vs Control 2

| Activation |  |  |  |  |  |
| --- | --- | --- | --- | --- | --- |
| | $V_{50}$ | $k$ | $V_{rev}$ | Peak current density | $n$ |
| Na<sup>+</sup> currents | mV | mV | mV | pA/pF |  |
| Control 1 | -29 ± 1 | 3 ± 1 | 17 ± 1 | -27 ± 1* | 14 |
| Control 2 | -34 ± 1 | 4 ± 1 | 15 ± 1 | -38 ± 1 | 15 |
| Ca<sup>2+</sup> currents |  |  |  |  |  |
| Control 1 | - 8 ± 1 | 8 ± 1 | 54 ± 1 | -10 ± 1 | 9 |
| Control 2 | -11 ± 1 | 8 ± 1 | 55 ± 2 | -10 ± 1 | 11 |

Activation parameters were calculated by data fitting to Boltzmann functions. $V_{50}$ is the voltage for half-maximal activation, $k$ is the slope factor and $n$ the number of cells. One-way ANOVA followed by Dunnett's multiple comparisons test. Values are expressed as mean ± s.e.m. \* $P < 0.05$ .

**Supplemental Table 7.**
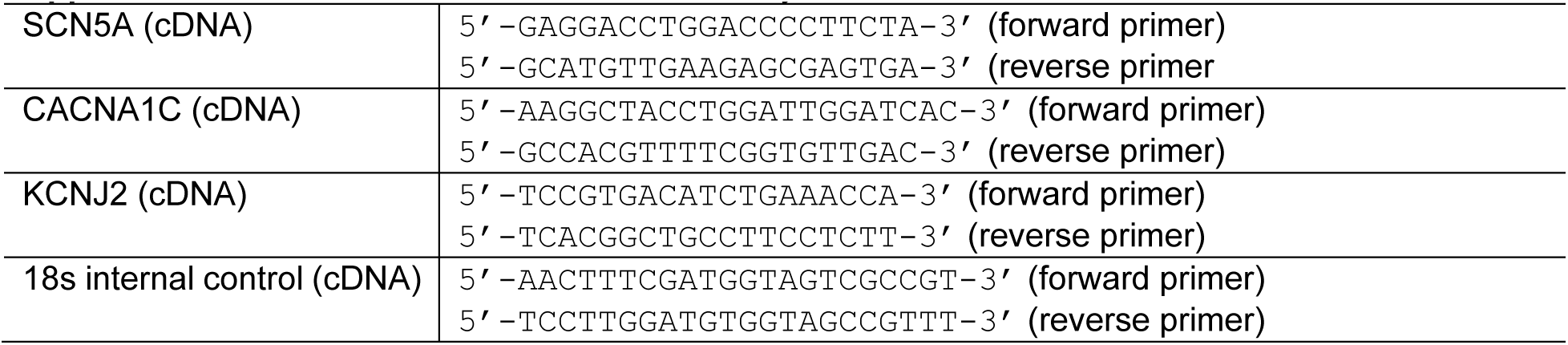
Primers used in mRNA analysis

## SUPPLEMENTAL METHODS

## 1. Ethics statement

We obtained skin biopsies from 2 hemizygous DMD patients, 1 heterozygous female, and a healthy patient after written informed consent in accordance with the Helsinki Committee for Experiments on Human Subjects at Sheba Medical Center, Ramat Gan, Israel (Approval number: 7603-09-SMC), and with IRB HUM00030934 approved by the University of Michigan Human IRB Committee. The use of iPS cells and iPSC-CMs was approved by the Human Pluripotent Stem Cell Research Oversight (HPSCRO, #1062) Committee of the University of Michigan and by the Spanish National Center for Cardiovascular Research (CNIC) Ethics Committee and the Regional Government of Madrid.

## 2. Generation of iPSCs

Cell lines were generated using Sendai virus CytoTune-iPS 2.0 Sendai reprogramming kit (Thermo Fisher) for transfection of Yamanaka’s factors: OCT4, KLF4, c-Myc, and SOX2, as described.^1, 2^ Subsequently, iPSCs were cultured on Matrigel (Corning)-coated 6-well plates with mTeSR1 medium (Stemcell Technologies) at 37°C with 5% CO_2_. iPSCs were passaged every 5 days at a ratio of 1:6 by mechanical dissociation using 1 mL/well of Versene solution (Invitrogen) following incubation at 37 °C for 7 min. DMD iPSCs were transported from Israel in dry ice to Michigan and to CNIC where they were differentiated to iPSC-CMs and used for the initial (Michigan) and syntrophin rescue (CNIC) studies. All iPSCs were tested for pluripotency before starting cardiomyocyte differentiation protocols. All of cells correlated well with the expression status of the pluripotency factors. Differentiation markers were also assessed.

## 3. Patient-specific iPSC-CMs monolayers: Differentiation into cardiomyocytes, adapted from ^3^

iPSC-CMs were generated by directed differentiation, modulating Wnt/β-catenin signaling.^4^ Briefly, iPSCs were cultured for 5–6 days on Matrigel-coated (Corning, 100 μg/mL) 6-well plates in StemMACs iPSC Brew XF medium (Miltenyi Biotec). Then, iPSCs were dissociated using 1 mL/well Versene solution at 37°C for 7 min and reseeded as monolayers on Matrigel-coated 12-well plates at a density of 8.5 × 10^6^ cells/well in StemMACs iPSC Brew XF medium supplemented with 5 μmol/L ROCK inhibitor (Miltenyi); medium was replaced every day. After 2 days, when monolayers reached 100% confluence, the medium was changed to RPMI supplemented with B27 minus insulin (Invitrogen) containing 10 μmol/L CHIR99021; this day was labelled as day 1 of differentiation. On day 2, the medium was changed to RPMI supplemented with B27 minus insulin. On day 4, the medium was changed to RPMI supplemented with B27 minus insulin, containing 10 μmol/L of IWP-4. On day 6, the medium was changed to RPMI supplemented with B27 minus insulin. Finally, from the 8^th^ day onwards, the medium was changed to RPMI supplemented with B27 complete supplement, RPMI +B27 media (Invitrogen).

## 4. Patient-specific iPSC-CMs monolayers: Post directed differentiation iPSC-CMs purification using MACs negative selection

The directed differentiation method used here does not generate a completely pure iPSC-CM population. Hence, the following purification steps preceded any characterization or experiments. iPSC-CMs ≥30 days in culture were washed with DPBS (Gibco) and dissociated using 1 mL of 0.25% Trypsin/EDTA per well. Next, 2 mL of EB20 media was added per well of dissociated cells, each well was triturated and then transferred into a sterile 15-mL conical. The EB20 media was composed of: 80% DMEM/F12 (Gibco), 0.1 mM Non-Essential Amino Acids (Gibco), 1 mM L-Glutamine (Gibco), 0.1 mM β-mercaptoethanol (Gibco), 20% Fetal Bovine Serum (FBS, Corning), and 10 µM Blebbistatin (Toronto Research Chemicals). Collected cells were centrifuged at 900 RPM for 5 min at 4°C. *Purification*:^5, 6^ After removal of the supernatant, 6 mL of MACs Buffer was added followed by trituration. Cells were centrifuged again at 900 RPM for 5 min at 4°C. The supernatant was aspirated and 80 µL of MACs Buffer was added to resuspend the pellet. Then, 20 µL of non-cardiomyocyte depletion cocktail (-Biotin conjugated) primary antibody was added, flicked 5 times to mix and incubated on ice for 5 min. After primary antibody incubation, 1 mL MACs Buffer was added, and cells were gently triturated followed by a 900 RPM spin for 5 min at 4°C. The excess primary antibody was aspirated and 80 µL of MACs Buffer was used to resuspend the pellet. Next, it was mixed with 20 µL of anti-Biotin magnetic microbeads (secondary antibody) and incubated on ice for 5 min. In the meantime, LS columns with 30 µm separation filters were placed onto a Quadro MACS Separator magnet, 15-mL conical tubes were appropriately labeled and positioned under each column, and 3 mL of MACs Buffer was run through each column to prime for addition of cell suspension. After secondary antibody incubation, cells were mixed with 1 mL of MACs Buffer. Then, the cell suspension was added to the separating filter on top of the flowing column, followed by 3 × 3 mL of cold MACs Buffer washes while continuously collecting the total flow through. The flow through or iPSC-CMs fraction was triturated and 1 mL of the total suspension was placed in a 1.5-mL Eppendorf tube to count the iPSC-CMs using a Millipore Scepter with Sensor tips (60 µm), this 1 mL was added back to the iPSC-CMs suspension total. Next, the purified (98-99%) iPSC-CMs were centrifuged, the supernatant aspirated, and then resuspended in media for plating. *Plating*. The purified iPSC-CMs fractions were resuspended in EB20 media with 5 µM of ROCK inhibitor to 200–300k cells/200–300 µL volume and plated as monolayers on 22 mm × 22 mm cut Matrigel-diluted in DMEM/F12 media) PDMS. The plate was transferred to the incubator at 37°C and 5% CO_2_ for 2 hours. Next, 3 mL of EB20/ROCK inhibitor media was added to each well. After 2 days, iPSC-CMs were washed with 3 mL DPBS with Ca^2+^ and Mg^2+^ (Gibco) followed by addition of 3 mL of RPMI +B27 media; media was changed every 3 days. The highly purified iPSC-CMs were in monolayer culture on Matrigel-PDMS for at least 7 days after plating to induce maturation. Then monolayers were dissociated with 0.25% Trypsin/EDTA and re-plated onto Matrigel-coated micropatterned PDMS. All iPSC-CM selection materials were purchased from Miltenyi Biotec, except for culture media which was mixed in the laboratory. All the tests carried out in this study were performed using at least 3 separate cardiomyocyte differentiations.

## 5. Micropatterning on PDMS (adapted from ref^7^)

Micropatterned area was 1 cm × 1 cm total, each island was 100 µm length × 15 µm width and islands were spaced 80 µm from each other. *Preparing PDMS stamps.* The surface of stamps was cleaned with scotch tape followed by sonication in 70% ethanol/milli-Q water for at least 20 min. In a sterile hood, they were allowed to dry and then, incubated with 250 μL Matrigel (100 ☐g/mL) diluted in water at room temperature for at least 1 h. *Preparing PDMS substrates in 6-well plates.* 18 mm PDMS circles were sonicated in 70% ethanol for 20 min and transferred to a 6-well plate after shaking excess EtOH off. When ready for microprinting, the culture dish was UVO treated with the lid off for 9 min. *Microprinting.* While UVO is performed on PDMS circles, the Matrigel solution from the PDMS stamps was aspirated. After UVO was completed, dried stamps were inverted onto each PDMS circle and removed one by one after ∼2 min. Later, the micropatterned PDMS plate was incubated with pluronic-F127 overnight at room temperature. *Single cell re-plating.* Before re-plating iPSC-CMs, micropattern plates were cleaned with 3× PSA (Penicillin-Streptomycin-Amphotericin B solution; Thermo Scientific) diluted in PBS (Gibco) for 1 h, and exposed to UV light for 15 min. iPSC-CMs were dissociated from monolayers using trypsin 0.25% with EDTA for 8–10 min and adding RPMI media containing 10% FBS after dissociation. Next, dissociated iPSC-CMs were transferred through a 70 μm filter into a 50-mL conical tube. The iPSC-CM suspension was centrifuged at 700 RPM for 3 min. Subsequently, iPSC-CMs were re-suspended in warm RPMI/B27+ (with insulin) media supplemented with 2% FBS and 5 μM ROCK inhibitor (re-plating media). Finally, ∼30k iPSC-CMs in 350 μL re-plating media were placed in the center of the micropatterned area. After ∼5 h, 2 mL of re-plating media was added very gently. Plate was returned to the incubator and media change was performed at days 1 and 3 after re-plating. iPSC-CMs were on micropatterns at least 4 days prior to patch-clamping experiments.

## 6. Electrophysiology

Standard patch-clamp recording techniques were used to measure action potentials, *I*_Na_, *I*_CaL_, and *I*_K1 3, 8_. All the experiments were performed at room temperature (22°C–25°C), except for the AP that were recorded at 37°C.

Voltage-clamp experiments were controlled with a Multiclamp 700B amplifier and a Digidata 1440A acquisition system (Molecular Devices). Data were filtered at 5 kHz and sampled at 5–20 kHz. Activation curve data were fitted to a Boltzmann equation, of the form g = g_max_ / (1 + exp (V_50_ − V_m_) / *k*), where g is the conductance, g_max_ the maximum conductance, V_m_ is the membrane potential, V_50_ is the voltage at which half of the channels are activated, and *k* is the slope factor.

Pipettes were formed from aluminosilicate glass (AF150-100-10; Science Products) with a P-97 horizontal puller (Sutter Instruments), and had resistances between 2 and 3 MΩ for patch-clamp experiments and 5–7 MΩ for current-clamp recordings when filled with the respective pipette solutions (see below).

### Action potential recordings

APs were elicited at 1 and 2 Hz in current-clamp mode using a programmable digital stimulator. The iPSC-CMs were bathed in 148 mM NaCl, 0.4 mM NaH_2_PO_4_, 1 mM MgCl_2_, 5.4 mM KCl, 1.8 mM CaCl_2_, 15 mM HEPES, and 5.5 mM glucose, pH = 7.4 adjusted with NaOH. The pipette solution contained 150 mM KCl, 1 mM MgCl_2_, 1 mM EGTA, 5 mM HEPES, 5 mM phosphocreatine, 4.4 mM K_2_ATP, and 2 mM β-hydroxybutyric acid, pH = 7.2 adjusted with KOH. Action potential properties including, maximum diastolic potential, overshoot, action potential amplitude and action potential duration were analyzed using custom-made software developed by Krzysztof Grzeda for the Center of Arrhythmia Research, University of Michigan. Maximum upstroke velocity was estimated using OriginPro 9 (OriginLab Corporation). For current clamp experiments, we selected control iPSC-CMs with ventricular-like action potentials showing a rectangular configuration, upstroke velocities (dV/dtmax) ≥40 V/s and amplitudes of 100 mV in the controls. Cells with a triangulated action potential (i.e., atrial-like), depolarized MDP and steep phase-4 depolarization (node-like) were discarded. All iPSC-CMs selected for patch clamping were quiescent and required external stimulation to generate action potentials.

### Single currents

For *I*_Na_, we used a pulse protocol from -80 mV to +55 mV with a holding potential of -160 mV. Recordings were made in a bath solution that consisted of 10 mM NaCl, 1 mM MgCl_2_, 0.1 mM CdCl_2_, 20mM HEPES, 11 mM Glucose, 60 mM CsCl, and 72.5 mM Choline chloride, pH = 7.35 adjusted with CsOH. The pipette solution contained 60 mM CsF, 5 mM NaCl, 10 mM EGTA, 5 mM HEPES, 5 mM MgATP, and 75 mM Choline chloride, pH = 7.2 adjusted with CsOH.

*I*_K1_ was elicited from a holding potential of -50 mV by 500-ms steps from -120 to +40 mV. The external recording solution contained 148 mM NaCl, 0.4 mM NaH_2_PO4, 1 mM MgCl_2_, 5.5 mM Glucose, 1.8 mM CaCl_2_, 5.4 mM KCl, 15 mM HEPES, and 5 µM Nifedipine, pH = 7.4 adjusted with NaOH. 1 mM BaCl_2_ was used to isolate *I*_K1_ from other background currents (subtract solution). The internal solution contained 1 mM MgCl_2_, 5 mM EGTA, 140 mM KCl, 5 mM HEPES, 5 mM Phosphocreatine, 4.4 mM K_2_ATP, and 2 mM β-Hydroxybutyric acid, pH = 7.2 adjusted with KOH.

*I*_CaL_ was evoked applying a voltage-step protocol from -40 mV to +80 mV with a holding potential of -50 mV. The iPSC-CMs were bathed in 137 mM TEA-Cl, 5.4 mM CsCl, 1 mM MgCl_2_, 1.8 mM CaCl_2_, 4 mM Aminopyridine, 10 mM HEPES, 30 µM TTX, and 11 mM Glucose, pH = 7.4 adjusted with CsOH. The pipette solution contained 20 mM TEA-Cl, 120 mM CsCl, 1 mM MgCl_2•_6H_2_O, 5.2 mM Mg-ATP, 10 mM HEPES, and 10 mM EGTA, pH = 7.2 adjusted with CsOH.

Chemicals were purchased from Sigma. Data analysis was performed using pClamp 10.2 software package (Axon Instruments).

## 7. RT-PCR

For quantitative evaluation of the steady-state mRNA expression in iPSC-CM cultures, total RNA was prepared using the RNeasy Mini Kit (Qiagen), including DNAse treatment. 300 ng of RNA were reversed transcribed and converted to cDNA with oligo(dT)15 primers using reverse transcriptase according to manufacturer’s specifications, SuperScript III First-Strand Synthesis System (Invitrogen). Quantitative PCR was performed using Sybergreen Master Mix (Applied Biosystems) in the presence of sense- and antisense-primers (10 µM) for *SCN5A*, *CACNA1C* and *KCNJ2*, as described previously (Table S3)._9_ The PCR condition consisted of 95°C for 5 min, followed by 40 cycles of 95°C for 15 secs and 60°C for 1 min, followed by melting-curve analysis to verify the correctness of the amplicon.

The samples were analyzed in biological triplicates using the primers listed in supplemental table 1 and run in a StepOnePlus Real-Time PCR system (Applied Biosystems). The expression of the mRNA of the gene of interest relative to the internal control 18s rRNA in samples from control, hemizygous and heterozygous iPSC-CMs was calculated by the ΔΔCT method, based on the threshold cycle (CT), as fold change = 2^−(ΔΔCT), where ΔCT = CT_gene of interest_ − CT_18S_ and ΔΔCT = ΔCT_hemizygous/_*_heterozygous_* _iPSC-CMs_ − ΔCT_control iPSC-CMs 10_. From each experiment, the cDNA of 3 cell culture wells were measured as biological replicates of each cell line. Each cell culture well was measured from at least 3 separate cardiomyocyte differentiation cultures as technical replicates.

## 8. Wester Blotting: Cell surface protein biotinylation/Western Blot

iPSC-CMs were plated as above, and membrane proteins were biotinylated. iPSC-CMs monolayers were washed twice with ice cold PBS and biotinylated for 1 h at 4°C using PBS containing 1.5 mg of EZ Link Sulfo-NHS-SS-Biotin (Thermo Scientific). Next, each monolayer was washed 3× with PBS before and after 10 min/4°C incubation with PBS/100 mM Glycine (to quench unlinked biotin). Finally, iPSC-CMs were lysed for 1 h at 4°C with lysis buffer containing (in mM, pH = 7.4): 150 NaCl, 25 Tris, 1% Triton X, and 1% Sodium deoxycholate, supplemented with protease inhibitors consisting of 1 µg/mL Benzamidine, 2 µg/mL Leupeptin, and 2 µg/mL Pepstatin A.

## 9. Wester Blotting: Protein precipitation

Pull-down experiments were conducted overnight at 4°C with 30 µg of biotinylated protein dissolved in 100 μL of lysis buffer and 30 μL of Pierce Streptavidin magnetic beads (Thermo Scientific). Next day, magnetic beads were washed three times with lysis buffer, and the first supernatant was collected. 25 µL of 4× loading buffer were then added to the magnetic beads. Before loading samples into the gel, they were heated at 50°C for 5 min.

## 10. SDS/PAGE and immunoblotting

Proteins were resolved in 4–20% SDS-PAGE gels and transferred to iBlot^®^ stacks with regular PVDF membranes using the Life Technologies iBlot2 system. Nonspecific binding sites were blocked with 5% albumin in PBS-T (in mM, 3 KH_2_PO_4_, 10 Na_2_HPO_4_, 150 NaCl, and 0.1% Tween 20, pH = 7.2–7.4) for 30 min at room temperature. Membranes were probed with the anti-human Na_V_1.5 or Kir2.1 antibody diluted in 5% albumin/PBS-T overnight at 4°C. After washing 3×/10 min, membranes were incubated for 1 h with a secondary horseradish peroxidase-conjugated antibody diluted in 5% albumin/PBS-T. Subsequently, membranes were washed 3×/10 min with PBS-T. Signals were detected with the SuperSignal West Pico Chemiluminescent substrate (Thermo Scientific). Expression of Na_V_1.5 and Kir2.1 was quantified using Image Lab software (Bio-Rad).

Primary antibodies were prepared in block solution. Mouse anti-Cardiac Troponin T antibody (1:1000, #Ab10214, Abcam) was used to identify cTnT as the marker for cardiomyocytes. Rabbit anti-Na_V_1.5 antibody (clone ASC-013, Alomone Labs) was used for Na_V_1.5 protein expression (1:500), mouse anti-Kir2.1 antibody (clone N112B/14, University of California at Davis/Nacional Institutes of Health 105 NeuroMab Facility) was used for Kir2.1 protein expression (1:500), mouse anti-Dystrophin (1:000, #D8043, Sigma) was used to detect the Dp427 dystrophin isoform. Mouse anti-Actinin antibody (1:1000, #A7811, Sigma) was used to detect Actinin, loading control in total protein analysis. A mouse antibody (#Ab7671, Abcam) was used to detect the Na-KATPase, positive control for biotinylation assays. Rabbit anti-Connexin antibody (1:1000, #C6219, Sigma) was used to detect Connexin 43. HRP-conjugated secondary antibodies (mouse HRP #115-035-146 and rabbit HRP #111-035-144) were obtained from Jackson ImmunoResearch Laboratories for Western blot analysis.

## 11. Immunofluorescence

iPSC-CMs were seeded on micropatterned Matrigel-coated 6-well plates and fixed with 2% paraformaldehyde/PBS for 15 min. Cells were incubated for 10 min at a 1:100 dilution of wheat germ agglutinin (WGA) Alexa 488 (ThermoScientific), washed with PBS, and re-fixed in 4% formaldehyde in PBS at RT. Then, hiPSC-CMs were washed 5 min with PBS and blocked with block solution (PBS + 5% BSA + 0.4% Triton X) for 1 h. Incubation with primary antibodies was done in block solution for 1.5 h in a humidity chamber. To washout the excess of primary antibody, hiPSC-CMs were washed 3×/5min with PBS. Next, secondary antibodies in block solution were added to each slip and incubated for 1 h in a humidity chamber at room temperature. hiPSC-CMs were kept in dark, washed with PBS 3×/5 min, and mounted with PermaFluor Aqueous (Thermo Fisher) and coverslip.

Primary antibodies were used at different dilutions in block solution: Troponin I (#MAB1691, Millipore) was used at 1:500, Kir2.1 (#APC-026, Alomone) antibody was used at 1:200, Nav1.5 (#AGP-008, Alomone) was used at 1:200, Dystrophin MANDRA1 (#D8043, Sigma) was used at 1:100, and Phalloidin 488 (#A12379, Invitrogen) at 1:500 (it comes with a fluorophore conjugated so no secondary Ab incubation was needed, stains F-actin). Secondary Ab for cTnI was Cy3 Goat anti-Mouse IgG (1:400, #115-167-003, Jackson Immuno Research), for anti-dystrophin MANDRA1, the Cy3 Rat anti-mouse IgG (1:200, #415-165-166, Jackson ImmunoResearch) was used, for Kir2.1 was used Alexa Fluor 568 Goat Anti Rabbit IgG (H+L) (Invitrogen, 1:500) and for Nav1.5 was used Alexa Fluor 680 Goat Anti Guinea Pig IgG (H+L) (Invitrogen, 1:500). Both secondary Abs were diluted in block solution containing 1:10,000 DAPI (#D9542, Sigma) stain dilution. Immunostained preparations were analyzed by confocal microscopy, using a Nikon A1R confocal microscope 102 (Nikon Instruments Inc) Leica SP8 confocal microscope (Leica Microsystems) to determine protein localization.

## 12. Optical Mapping

iPSC-CMs were plated as monolayers at a density of ∼50k iPSC-CMs in RPMI/B27+ media. After 7 days in culture, media was removed and each iPSC-CMs monolayer was washed with Hank’s balanced salt solution with Ca^2+^ and Mg^2+^ added (HBSS^++^, Thermo Scientific) to remove remaining media. Next, iPSC-CMs were incubated with the FluoVolt membrane potential probe (F10488; Thermo Scientific) diluted in HBSS^++^, as reported before ^11^. After a 30-minute incubation time, iPSC-CMs were washed with HBSS^++^ and then heated at 35°C before optical mapping recordings. All iPSC-CMs monolayers displayed pacemaker activity, and the spontaneous and paced APs were recorded using a charge-coupled device camera (200 fps, 80 × 80 pixels; Red-Shirt Little Joe) with the appropriate emission filters and light-emitting diode illumination ^12^. The recorded videos were filtered in both the time and the space domain, and CV was measured as described previously ^3, 13^.

## 13. Generation and Stable Transfection of *SNTA1-IRES-GFP* using PiggyBac Transposon Integration Methods

Non-viral piggy-bac vector (1 µg) encoding SNTA1-IRES-GFP were co-transfected with mouse transposase-expression vector (250 ng) by electroporation (Amaxa® 4D-Nucleofector, Lonza) into iPSCs cells (∼1.10^6^ cells/electroporation). After 3-5 days GFP positive cells were selected by FACS sorter (BD FACSAria Cell Sorter, BD BioSciences) and grow-up. Every week, until three times, fluorescence was confirmed, and cells sorted to confirm cDNA stable integration into the cells. After that, iPSC-CMs differentiation protocol was applied as stated above.

## 14. Statistics

Statistical analyses were performed with Prism 8 (GraphPad Software). Values were first tested for normality (Shapiro-Wilk test) before statistical evaluation. Nonparametric Mann-Whitney rank test (two-tailed) was used. Multiple comparisons were analyzed using two-way analysis of variance (ANOVA) followed by Sidak’s test. All data are shown as mean ± s.e.m. P < 0.05 (2-tailed) was considered significant. Unless stated otherwise, the number n of observations indicated reflects the number of iPSC-CMs recorded from each cell line from at least 3 differentiations.

**Supplemental video 1.** Focal discharges in Female iPSC-CMs monolayer.

**Supplemental video 2.** Local conduction block in Female iPSC-CMs monolayer.

## Notes

### Competing Interest Statement

The authors have declared no competing interest.

## References

1. Hoffman EP, Monaco AP, Feener CC, Kunkel LM. Conservation of the Duchenne muscular dystrophy gene in mice and humans. Science 1987;238:347–350.

2. Anderson JL, Head SI, Rae C, Morley JW. Brain function in Duchenne muscular dystrophy. Brain 2002;125:4–13.

3. Corrado G, Lissoni A, Beretta S, Terenghi L, Tadeo G, Foglia-Manzillo G, Tagliagambe LM, Spata M, Santarone M. Prognostic value of electrocardiograms, ventricular late potentials, ventricular arrhythmias, and left ventricular systolic dysfunction in patients with Duchenne muscular dystrophy. Am J Cardiol 2002;89:838–841.

4. Finsterer J, Stollberger C, Freudenthaler B, Simoni D, Hoftberger R, Wagner K. Muscular and cardiac manifestations in a Duchenne-carrier harboring a dystrophin deletion of exons 12-29. Intractable Rare Dis Res 2018;7:120–125.

5. Shirokova N, Niggli E. Cardiac phenotype of Duchenne Muscular Dystrophy: insights from cellular studies. J Mol Cell Cardiol 2013;58:217–224.

6. Yilmaz A, Sechtem U. Cardiac involvement in muscular dystrophy: advances in diagnosis and therapy. Heart 2012;98:420–429.

7. Yilmaz A, Gdynia HJ, Mahrholdt H, Sechtem U. Cardiovascular magnetic resonance reveals similar damage to the heart of patients with Becker and limb-girdle muscular dystrophy but no cardiac symptoms. J Magn Reson Imaging 2009;30:876–877.

8. Petrof BJ, Shrager JB, Stedman HH, Kelly AM, Sweeney HL. Dystrophin protects the sarcolemma from stresses developed during muscle contraction. Proc Natl Acad Sci U S A 1993;90:3710–3714.

9. Constantin B. Dystrophin complex functions as a scaffold for signalling proteins. Biochimica et biophysica acta 2014;1838:635–642.

10. Gavillet B, Rougier JS, Domenighetti AA, Behar R, Boixel C, Ruchat P, Lehr HA, Pedrazzini T, Abriel H. Cardiac sodium channel Nav1.5 is regulated by a multiprotein complex composed of syntrophins and dystrophin. Circ Res 2006;99:407–414.

11. Milstein ML, Musa H, Balbuena DP, Anumonwo JM, Auerbach DS, Furspan PB, Hou L, Hu B, Schumacher SM, Vaidyanathan R, Martens JR, Jalife J. Dynamic reciprocity of sodium and potassium channel expression in a macromolecular complex controls cardiac excitability and arrhythmia. Proc Natl Acad Sci U S A 2012;109:E2134–2143.

12. Lohan J, Culligan K, Ohlendieck K. Deficiency in Cardiac Dystrophin Affects the Abundance of the $\alpha$ -/ $\beta$ -Dystroglycan Complex. J Biomed Biotechnol 2005;2005:28–36.

13. Koenig X, Rubi L, Obermair GJ, Cervenka R, Dang XB, Lukacs P, Kummer S, Bittner RE, Kubista H, Todt H, Hilber K. Enhanced currents through L-type calcium channels in cardiomyocytes disturb the electrophysiology of the dystrophic heart. Am J Physiol Heart Circ Physiol 2014;306:H564–573.

14. Rubi L, Koenig X, Kubista H, Todt H, Hilber K. Decreased inward rectifier potassium current IK1 in dystrophin-deficient ventricular cardiomyocytes. Channels (Austin) 2017;11:101–108.

15. Koenig X, Dysek S, Kimbacher S, Mike AK, Cervenka R, Lukacs P, Nagl K, Dang XB, Todt H, Bittner RE, Hilber K. Voltage-gated ion channel dysfunction precedes cardiomyopathy development in the dystrophic heart. PLoS One 2011;6:e20300.

16. Albesa M, Ogrodnik J, Rougier JS, Abriel H. Regulation of the cardiac sodium channel Nav1.5 by utrophin in dystrophin-deficient mice. Cardiovasc Res 2011;89:320–328.

17. Petitprez S, Zmoos AF, Ogrodnik J, Balse E, Raad N, El-Haou S, Albesa M, Bittihn P, Luther S, Lehnart SE, Hatem SN, Coulombe A, Abriel H. SAP97 and dystrophin macromolecular complexes determine two pools of cardiac sodium channels Nav1.5 in cardiomyocytes. Circ Res 2011;108:294–304.

18. Leonoudakis D, Conti LR, Anderson S, Radeke CM, McGuire LM, Adams ME, Froehner SC, Yates JR, 3rd, Vandenberg CA. Protein trafficking and anchoring complexes revealed by proteomic analysis of inward rectifier potassium channel (Kir2.x)-associated proteins. J Biol Chem 2004;279:22331-22346.

19. Matamoros M, Perez-Hernandez M, Guerrero-Serna G, Amoros I, Barana A, Nunez M, Ponce-Balbuena D, Sacristan S, Gomez R, Tamargo J, Caballero R, Jalife J, Delpon E. Nav1.5 N-terminal domain binding to alpha1-syntrophin increases membrane density of human Kir2.1, Kir2.2 and Nav1.5 channels. Cardiovasc Res 2016;110:279–290.

20. Ponce-Balbuena D, Guerrero-Serna G, Valdivia CR, Caballero R, Diez-Guerra FJ, Jimenez-Vazquez EN, Ramirez RJ, Monteiro da Rocha A, Herron TJ, Campbell KF, Willis BC, Alvarado FJ, Zarzoso M, Kaur K, Perez-Hernandez M, Matamoros M, Valdivia HH, Delpon E, Jalife J. Cardiac Kir2.1 and NaV1.5 Channels Traffic Together to the Sarcolemma to Control Excitability. Circ Res 2018;122:1501–1516.

21. Perez-Hernandez M, Matamoros M, Alfayate S, Nieto-Marin P, Utrilla RG, Tinaquero D, de Andres R, Crespo T, Ponce-Balbuena D, Willis BC, Jimenez-Vazquez EN, Guerrero-Serna G, da Rocha AM, Campbell K, Herron TJ, Diez-Guerra FJ, Tamargo J, Jalife J, Caballero R, Delpon E. Brugada syndrome trafficking-defective Nav1.5 channels can trap cardiac Kir2.1/2.2 channels. JCI Insight 2018;3.

22. Eisen B, Ben Jehuda R, Cuttitta AJ, Mekies LN, Reiter I, Ramchandren S, Arad M, Michele DE, Binah O. Generation of Duchenne muscular dystrophy patient-specific induced pluripotent stem cell line lacking exons 45-50 of the dystrophin gene (IITi001-A). Stem Cell Res 2018;29:111–114.

23. Eisen B, Ben Jehuda R, Cuttitta AJ, Mekies LN, Shemer Y, Baskin P, Reiter I, Willi L, Freimark D, Gherghiceanu M, Monserrat L, Scherr M, Hilfiker-Kleiner D, Arad M, Michele DE, Binah O. Electrophysiological abnormalities in induced pluripotent stem cell-derived cardiomyocytes generated from Duchenne muscular dystrophy patients. J Cell Mol Med 2019;23:2125–2135.

24. Herron TJ, Rocha AM, Campbell KF, Ponce-Balbuena D, Willis BC, Guerrero-Serna G, Liu Q, Klos M, Musa H, Zarzoso M, Bizy A, Furness J, Anumonwo J, Mironov S, Jalife J. Extracellular Matrix-Mediated Maturation of Human Pluripotent Stem Cell-Derived Cardiac Monolayer Structure and Electrophysiological Function. Circ Arrhythm Electrophysiol 2016;9:e003638.

25. Kuo PL, Lee H, Bray MA, Geisse NA, Huang YT, Adams WJ, Sheehy SP, Parker KK. Myocyte shape regulates lateral registry of sarcomeres and contractility. Am J Pathol 2012;181:2030–2037.

26. da Rocha AM, Campbell K, Mironov S, Jiang J, Mundada L, Guerrero-Serna G, Jalife J, Herron TJ. hiPSC-CM Monolayer Maturation State Determines Drug Responsiveness in High Throughput Pro-Arrhythmia Screen. Sci Rep 2017;7:13834.

27. de Souza F, Bittar Braune C, Dos Santos Nucera APC. Duchenne muscular dystrophy: an overview to the cardiologist. Expert Rev Cardiovasc Ther 2020;18:867–872.

28. Szabo SM, Salhany RM, Deighton A, Harwood M, Mah J, Gooch KL. The clinical course of Duchenne muscular dystrophy in the corticosteroid treatment era: a systematic literature review. Orphanet J Rare Dis 2021;16:237.

29. Ribeiro AJ, Ang YS, Fu JD, Rivas RN, Mohamed TM, Higgs GC, Srivastava D, Pruitt BL. Contractility of single cardiomyocytes differentiated from pluripotent stem cells depends on physiological shape and substrate stiffness. Proc Natl Acad Sci U S A 2015;112:12705–12710.

30. Taggart P, Sutton PM, Boyett MR, Lab M, Swanton H. Human ventricular action potential duration during short and long cycles. Rapid modulation by ischemia. Circulation 1996;94:2526–2534.

31. Grandi E, Pandit SV, Voigt N, Workman AJ, Dobrev D, Jalife J, Bers DM. Human atrial action potential and Ca2+ model: sinus rhythm and chronic atrial fibrillation. Circ Res 2011;109:1055–1066.

32. Fayssoil A, Nardi O, Orlikowski D, Annane D. Cardiomyopathy in Duchenne muscular dystrophy: pathogenesis and therapeutics. Heart Fail Rev 2010;15:103–107.

33. Finsterer J, Stollberger C. The heart in human dystrophinopathies. Cardiology 2003;99:1–19.

34. Abriel H. Roles and regulation of the cardiac sodium channel Na v 1.5: recent insights from experimental studies. Cardiovasc Res 2007;76:381–389.

35. Yotsukura M, Miyagawa M, Tsuya T, Ishihara T, Ishikawa K. A 10-year follow-up study by orthogonal Frank lead ECG on patients with progressive muscular dystrophy of the Duchenne type. J Electrocardiol 1992;25:345–353.

36. Perloff JK. Cardiac rhythm and conduction in Duchenne’s muscular dystrophy: a prospective study of 20 patients. J Am Coll Cardiol 1984;3:1263–1268.

37. Viola HM, Davies SM, Filipovska A, Hool LC. L-type Ca(2+) channel contributes to alterations in mitochondrial calcium handling in the mdx ventricular myocyte. Am J Physiol Heart Circ Physiol 2013;304:H767–775.

38. Araishi K, Sasaoka T, Imamura M, Noguchi S, Hama H, Wakabayashi E, Yoshida M, Hori T, Ozawa E. Loss of the sarcoglycan complex and sarcospan leads to muscular dystrophy in beta-sarcoglycan-deficient mice. Hum Mol Genet 1999;8:1589–1598.

39. Gee SH, Madhavan R, Levinson SR, Caldwell JH, Sealock R, Froehner SC. Interaction of muscle and brain sodium channels with multiple members of the syntrophin family of dystrophin-associated proteins. J Neurosci 1998;18:128–137.

40. Hara H, Niwano S, Ito H, Karakawa M, Ako J. Evaluation of R-wave offset in the left chest leads for estimating the left ventricular activation delay: An evaluation based on coronary sinus electrograms and the 12-lead electrocardiogram. J Electrocardiol 2016;49:148–153.

41. Villa CR, Czosek RJ, Ahmed H, Khoury PR, Anderson JB, Knilans TK, Jefferies JL, Wong B, Spar DS. Ambulatory Monitoring and Arrhythmic Outcomes in Pediatric and Adolescent Patients With Duchenne Muscular Dystrophy. Journal of the American Heart Association 2015;5.

42. Ronaldson-Bouchard K, Yeager K, Teles D, Chen T, Ma S, Song L, Morikawa K, Wobma HM, Vasciaveo A, Ruiz EC, Yazawa M, Vunjak-Novakovic G. Engineering of human cardiac muscle electromechanically matured to an adult-like phenotype. Nat Protoc 2019;14:2781–2817.

43. Berecki G, Wilders R, de Jonge B, van Ginneken AC, Verkerk AO. Re-evaluation of the action potential upstroke velocity as a measure of the Na+ current in cardiac myocytes at physiological conditions. PLoS One 2010;5:e15772.

44. Sanford JL, Edwards JD, Mays TA, Gong B, Merriam AP, Rafael-Fortney JA. Claudin-5 localizes to the lateral membranes of cardiomyocytes and is altered in utrophin/dystrophin-deficient cardiomyopathic mice. J Mol Cell Cardiol 2005;38:323–332.

45. Okin PM, Devereux RB, Howard BV, Fabsitz RR, Lee ET, Welty TK. Assessment of QT interval and QT dispersion for prediction of all-cause and cardiovascular mortality in American Indians: The Strong Heart Study. Circulation 2000;101:61–66.

46. Shaw RM, Rudy Y. Electrophysiologic effects of acute myocardial ischemia: a theoretical study of altered cell excitability and action potential duration. Cardiovasc Res 1997;35:256–272.

47. Rougier JS, Gavillet B, Abriel H. Proteasome inhibitor (MG132) rescues Nav1.5 protein content and the cardiac sodium current in dystrophin-deficient mdx (5cv) mice. Front Physiol 2013;4:51.

48. Florian A, Rosch S, Bietenbeck M, Engelen M, Stypmann J, Waltenberger J, Sechtem U, Yilmaz A. Cardiac involvement in female Duchenne and Becker muscular dystrophy carriers in comparison to their first-degree male relatives: a comparative cardiovascular magnetic resonance study. Eur Heart J Cardiovasc Imaging 2016;17:326–333.

49. Holloway SM, Wilcox DE, Wilcox A, Dean JC, Berg JN, Goudie DR, Denvir MA, Porteous ME. Life expectancy and death from cardiomyopathy amongst carriers of Duchenne and Becker muscular dystrophy in Scotland. Heart 2008;94:633–636.

50. Kane AM, DeFrancesco TC, Boyle MC, Malarkey DE, Ritchey JW, Atkins CE, Cullen JM, Kornegay JN, Keene BW. Cardiac structure and function in female carriers of a canine model of Duchenne muscular dystrophy. Res Vet Sci 2013;94:610–617.

51. McCaffrey T, Guglieri M, Murphy AP, Bushby K, Johnson A, Bourke JP. Cardiac involvement in female carriers of duchenne or becker muscular dystrophy. Muscle Nerve 2017;55:810–818.

52. Richards DA, Byth K, Ross DL, Uther JB. What is the best predictor of spontaneous ventricular tachycardia and sudden death after myocardial infarction? Circulation 1991;83:756–763.

53. Madej-Pilarczyk A. Clinical aspects of Emery-Dreifuss muscular dystrophy. Nucleus 2018;9:268–274.

54. Lang SM, Shugh S, Mazur W, Sticka JJ, Rattan MS, Jefferies JL, Taylor MD. Myocardial Fibrosis and Left Ventricular Dysfunction in Duchenne Muscular Dystrophy Carriers Using Cardiac Magnetic Resonance Imaging. Pediatr Cardiol 2015;36:1495–1501.

## References

1. Eisen B, Ben Jehuda R, Cuttitta AJ, Mekies LN, Reiter I, Ramchandren S, Arad M, Michele DE and Binah O. Generation of Duchenne muscular dystrophy patient-specific induced pluripotent stem cell line lacking exons 45-50 of the dystrophin gene (IITi001-A). Stem Cell Res. 2018;29:111–114.

2. Eisen B, Ben Jehuda R, Cuttitta AJ, Mekies LN, Shemer Y, Baskin P, Reiter I, Willi L, Freimark D, Gherghiceanu M, Monserrat L, Scherr M, Hilfiker-Kleiner D, Arad M, Michele DE and Binah O. Electrophysiological abnormalities in induced pluripotent stem cell-derived cardiomyocytes generated from Duchenne muscular dystrophy patients. J Cell Mol Med. 2019;23:2125–2135.

3. Herron TJ, Rocha AM, Campbell KF, Ponce-Balbuena D, Willis BC, Guerrero-Serna G, Liu Q, Klos M, Musa H, Zarzoso M, Bizy A, Furness J, Anumonwo J, Mironov S and Jalife J. Extracellular Matrix-Mediated Maturation of Human Pluripotent Stem Cell-Derived Cardiac Monolayer Structure and Electrophysiological Function. Circ Arrhythm Electrophysiol. 2016;9:e003638.

4. Lian X, Zhang J, Azarin SM, Zhu K, Hazeltine LB, Bao X, Hsiao C, Kamp TJ and Palecek SP. Directed cardiomyocyte differentiation from human pluripotent stem cells by modulating Wnt/beta-catenin signaling under fully defined conditions. Nat Protoc. 2013;8:162–75.

5. Pekkanen-Mattila M, Hakli M, Polonen RP, Mansikkala T, Junnila A, Talvitie E, Koivisto JT, Kellomaki M and Aalto-Setala K. Polyethylene Terephthalate Textiles Enhance the Structural Maturation of Human Induced Pluripotent Stem Cell-Derived Cardiomyocytes. Materials (Basel). 2019;12.

6. Herron T, Monteiro da Rocha A and Campbell K. Cardiomyocyte purification from pluripotent stem cells. 2017.

7. Kuo PL, Lee H, Bray MA, Geisse NA, Huang YT, Adams WJ, Sheehy SP and Parker KK. Myocyte shape regulates lateral registry of sarcomeres and contractility. Am J Pathol. 2012;181:2030–7.

8. Caballero R, Utrilla RG, Amoros I, Matamoros M, Perez-Hernandez M, Tinaquero D, Alfayate S, Nieto-Marin P, Guerrero-Serna G, Liu QH, Ramos-Mondragon R, Ponce-Balbuena D, Herron T, Campbell KF, Filgueiras-Rama D, Peinado R, Lopez-Sendon JL, Jalife J, Delpon E and Tamargo J. Tbx20 controls the expression of the KCNH2 gene and of hERG channels. Proceedings of the National Academy of Sciences of the United States of America. 2017;114:E416–E425.

9. Bizy A, Guerrero-Serna G, Hu B, Ponce-Balbuena D, Willis BC, Zarzoso M, Ramirez RJ, Sener MF, Mundada LV, Klos M, Devaney EJ, Vikstrom KL, Herron TJ and Jalife J. Myosin light chain 2-based selection of human iPSC-derived early ventricular cardiac myocytes. Stem Cell Res. 2013;11:1335–47.

10. Schmittgen TD and Livak KJ. Analyzing real-time PCR data by the comparative C(T) method. Nat Protoc. 2008;3:1101–8.

11. da Rocha AM, Campbell K, Mironov S, Jiang J, Mundada L, Guerrero-Serna G, Jalife J and Herron TJ. hiPSC-CM Monolayer Maturation State Determines Drug Responsiveness in High Throughput Pro-Arrhythmia Screen. Sci Rep. 2017;7:13834.

12. Lee P, Bollensdorff C, Quinn TA, Wuskell JP, Loew LM and Kohl P. Single-sensor system for spatially resolved, continuous, and multiparametric optical mapping of cardiac tissue. Heart Rhythm. 2011;8:1482–91.

13. Campbell K, Calvo CJ, Mironov S, Herron T, Berenfeld O and Jalife J. Spatial gradients in action potential duration created by regional magnetofection of hERG are a substrate for wavebreak and turbulent propagation in cardiomyocyte monolayers. The Journal of physiology. 2012;590:6363–79.

